# Proteome-wide microarray-based screening of PAR-binding proteins

**DOI:** 10.1101/2022.06.06.494829

**Authors:** Bong Gu Kang, Sung-Ung Kang, Jae Jin Kim, Ji-Sun Kwon, Jean-Philippe Gagné, Seo Yun Lee, Soyeon Kim, Karl L. Sangwon, Shinwon Ha, Jun Seop Jeong, Yun-Il Lee, Heng Zhu, Dongsan Kim, Guy G. Poirier, Ho Chul Kang, Valina L. Dawson, Ted M. Dawson

## Abstract

Poly(ADP-ribose) (PAR) plays a crucial role in intracellular signaling and scaffolding through covalent modification or non-covalent binding to target proteins. The non-covalent binding PARylome has not been extensively characterized. Here we performed a PAR-binding screen using a human protein microarray that covers most of the human proteome to characterize the non-covalent binding PARylome. A total of 356 PAR-binding proteins were identified. The PAR-binding PARylome suggests that PAR-binding regulates a variety of biological processes beyond well-characterized DNA damage signaling and DNA repair. Proteins that may be reprogrammed by PAR-binding include signaling molecules, transcription factors, nucleic acid binding proteins, calcium binding proteins, ligases, oxidoreductases, enzymes, transferases, hydrolases, and receptors. The global database of PAR-binding proteins that we established will be a valuable tool for further in-depth analysis of the role of PARylation in a wide range of biological contexts.

## INTRODUCTION

Poly(ADP-ribosyl)ation (PARylation) is a post-translational protein modification that regulates a number of cellular processes including DNA repair and the maintenance of genome integrity, transcription, cell death and mRNA metabolism (Krishnakumar and Kraus, 2010; Luo and Kraus, 2012; Robert et al., 2013; Rouleau et al., 2010; Virag et al., 2013). Many of these processes are regulated by the covalent addition of poly(ADP-ribose) (PAR) – a biopolymer composed of linear or branched repeats of ADP-ribose – on target proteins via a class of enzymes called ADP-ribosyl transferases via a process designated PARylation (Kim et al., 2005; Luscher et al., 2021). In addition to its regulatory role as a covalent post-translational modification, PAR serves as a molecular scaffold that recruits other proteins, such as DNA damage response (DDR) factors, to regulate downstream signaling and repair pathways (Gibson and Kraus, 2012; Liu et al., 2017). In this scenario, proteins interact non-covalently with PAR to orchestrate the highly coordinated process of DNA damage repair. The binding of PAR via a non-covalent fashion (also referred to as PAR *reading*) is also a regulatory event in various other cellular pathways. PAR readers span several functions including the regulation of mitosis, cell death and chromatin modification, among others (Andrabi et al., 2006; Gibson and Kraus, 2012; Kim *et al*., 2005; Krishnakumar and Kraus, 2010; Yu et al., 2006).

Despite the crucial role PAR plays in numerous signaling pathways, the identification of PAR readers and the specific protein domains and motifs that confers affinity for PAR has not been extensively studied. This unusual functional duality of PAR has made it very challenging to analyze. The technology was limited in its ability to distinguish covalently PARylated substrates and their site-specific linkages from PAR readers. This complexity is reflected by the nature of the sugar–peptide bonds, the structural heterogeneity of PAR and the intricate network of interactions established between a constantly growing diversity of PAR readers with different binding affinities and binding properties to PAR.

Owing to the evolution of new instruments and methods, it is now possible to discriminate PARylated substrates from PAR readers. Specific dissociation techniques and spectral signatures were developed to preserve the labile post-translational ADP-ribosylation modifications via mass spectrometry (Jungmichel et al., 2013; Li and Chen, 2014; Luthi et al., 2021; Martello et al., 2013; Zhang et al., 2013). ADP-ribose probes and photoaffinity-based proteomics were specifically developed to assess PAR readers (Dasovich et al., 2021; Teloni and Altmeyer, 2016). Previously identified PAR readers include the PAR-binding motif (PBM), Macrodomain, PAR-binding Zinc-finger (PBZ), WWE, WD40, PIN, BRCT, Ob-fold, PIN, FHA domains, ZNFs, RNA recognition motifs (RRM), SR and KR repeats and intrinsically disordered regions (Altmeyer et al., 2015; Dasovich *et al*., 2021; Gagne et al., 2008; Gupte et al., 2017; Li and Chen, 2014; Teloni and Altmeyer, 2016). Despite these advances, the specific identification of PAR readers over PARylated substrates in complex biological samples has remained technically challenging.

Recently, high-density protein microarrays have been utilized to probe post-translational modifications and to gain insight into the repertoire of proteins that are regulated by specific modifications (Bertone and Snyder, 2005; Jones et al., 2006; Templin et al., 2002). Here, we used a high-density protein microarray containing 17,000 human proteins to define the human PAR-binding PARylome. Our approach allowed us to identify 356 proteins that binds PAR *in vitro* and to characterize novel PAR-binding motifs that add to the complexity of pathways involved in the cellular response to PARylation.

## RESULTS

### Identification of PAR-binding proteins using a 17K human protein microarray

To identify PAR-binding proteins (PAR readers), DHBB-purified PAR synthesized *in vitro* from PARP-1, was generated with biotin-labeled nicotinamide adenine dinucleotide (NAD^+^). PAR polymer was probed to a previously characterized high-density 17,000 (17K) human protein microarray (Hu et al., 2009; Lee et al., 2014) (Figure 1A).

**Figure 1.**
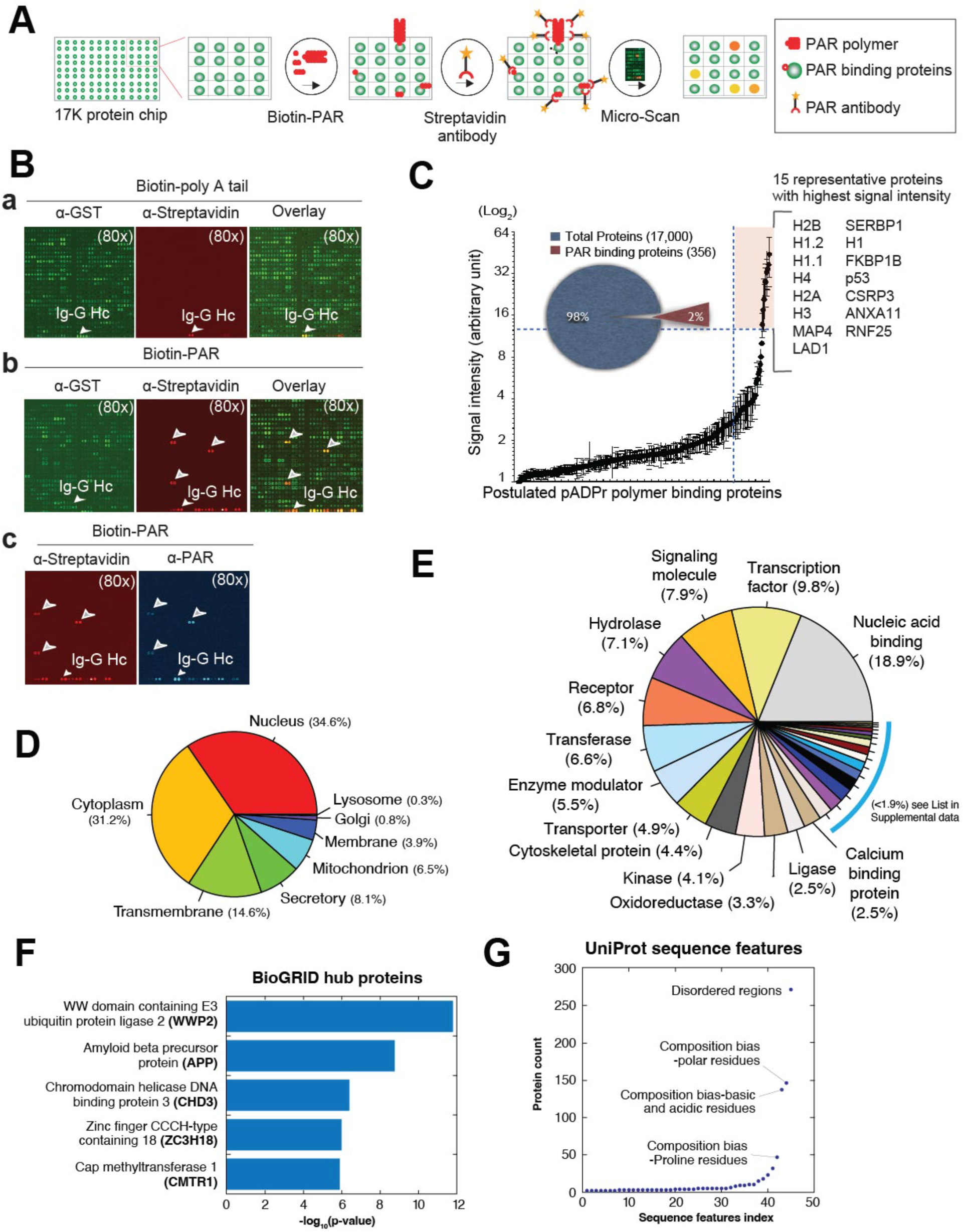
Identification and characterization of 356 unique PAR-binding proteins using 17k human protein microarray. (A) Workflow illustration of PAR-binding detection via protein microarray using biotin-PAR. (B) Representative scanned images of biotin-Poly A tails as (a) non-specific negative control background, (b) PAR-binding detection using anti-streptavidin antibodies, and (c) anti-PAR based detection for direct PAR-binding confirmation. (C) The 356 PAR-binding proteins were statistically filtered by signal intensity. The 15 representative proteins showing the highest affinity out of the 356 PAR-binding proteins are listed in the light orange box. (D) Subcellular localization of the 356 PAR-binding proteins. (E) Functional classification of the 356 PAR-binding proteins. (F) Hub proteins that connects to a high fraction of PAR-binding proteins based on BioGRID analysis (Biological General Repository for Interaction Datasets). (G) Classification of the most occurring sequence features found in PAR-binding proteins according to UniProt annotations.

Considering that PAR is a polyanionic polymer with a highly negative charge density and to exclude any non-specific ionic interactions with proteins, biotin-labeled polyadenylate [poly(A)] RNA was generated and probed against the protein microarray as a negative control (Figures 1B (panel a) and S1A). PAR-binding proteins were detected using anti-streptavidin antibodies that recognize the biotin label (Figures 1B (panel b) and S1B). We additionally confirmed direct PAR-binding by using an anti-PAR antibody that directly recognizes PAR (Figures 1B (panel c), S1C and S1D). Validation of PAR specificity was done via PAR glycohydrolase (PARG) treatment, which catabolizes PAR via hydrolysis of the ribose-ribose bond (Slade et al., 2011). PARG treatment completely erased the PAR-binding signal (Figures S1E and S1F). PAR-binding signal intensity was determined by a PAR/poly(A) normalization ratio and statistical filtering based on the signal intensity (Figure 1C and Table S1). 356 PAR-binding proteins were identified, which is approximately 2% of the proteins printed on the high-density microarray (Figure 1C and Table S1). Fifteen proteins, indicated by the points in the upper right shaded corner of the graph in Figure 1C, were observed to have the highest PAR-binding signal intensity (Figure 1C). As expected, known PAR-binding proteins such as histones (Panzeter et al., 1992; Reale et al., 2000; Realini and Althaus, 1992) and p53 (Fischbach et al., 2018; Kruger et al., 2019), were among the highest PAR binders. The specificity of the interaction was further underscored by the observation that positively charged lysine-rich histones do not bind measurably to negatively charged poly(A).

PAR readers were classified based on subcellular localizations, extensive protein interactions and the most occurring sequence features using multiple databases. PAR-binding proteins were primarily resident in the nucleus (34.6%), followed by the cytoplasm (31.2%) and transmembrane regions (14.6%) (Figure 1D and Table S2).

Functional classification of the PAR readers shows that the two most common protein functions were assigned to nucleic acid binding (18.9%) followed by transcription factor activity (9.8%), consistent with the role of PAR signaling in DNA damage repair and transcriptional regulation (Figure 1E). The data set of PAR readers (356 proteins) is significantly enriched in proteins that interacts with specific hub proteins, such as amyloid beta precursor protein (APP) and WW domain containing E3 ubiquitin protein ligase 2 (WWP2) with p-value of 1.67E-12 and 1.83E-09, respectively (Figure 1F and Table S3). 271 proteins (p-value of 8.72E-07) have disordered regions, consistent with the concept of PAR-seeded liquid demixing by intrinsically disordered proteins (Altmeyer *et al*., 2015) (Figure 1G and Table S4). Compositional bias in the amino acid sequences of PAR readers is consistent with the P/Q/E-rich motifs (Karlberg et al., 2013).

### Protein microarray detection of PAR-binding detection expands the PAR-binding proteome

68 PAR binding proteins identified via the protein microarray screen overlapped with two recent MS datasets of covalently PARylated proteins, generated using *in vitro* ADP-ribosylation coupled with quantitative proteomics or activated ion electron transfer dissociation (AI-ETD) (Buch-Larsen et al., 2020; Hendriks et al., 2021) (Figure 2A and Table S5). 42 PAR readers identified via the protein microarray screen overlapped with two affinity purification-based MS datasets of non-covalent PAR-binding proteins (Dasovich *et al*., 2021; Kliza et al., 2021). In total, 80 proteins identified as PAR readers via the protein microarray screen were previously found in non-covalent PAR binding or covalent PARylated protein datasets (Figure 2A and Table S5). Further comparisons with other strategies used to identify PAR readers and PARylated substrates including photoaffinity-based proteomics (Dasovich *et al*., 2021), GST-fusion macrodomain (Jungmichel *et al*., 2013), PAR antibody-based (Gagne *et al*., 2008) and boronate-affinity chromatography screens (Zhang *et al*., 2013) are highlighted in Figure 2B and Table S6. Domains that overlapped with the protein microarray PAR-binding and the MS-based datasets include WD40, BRCT, FHA, PIN, Ob-fold, ZNF (zf-C2H2) and RRM with the ZNF and RRM domains as the most common (Figure 2B and Table S6).

**Figure 2.**
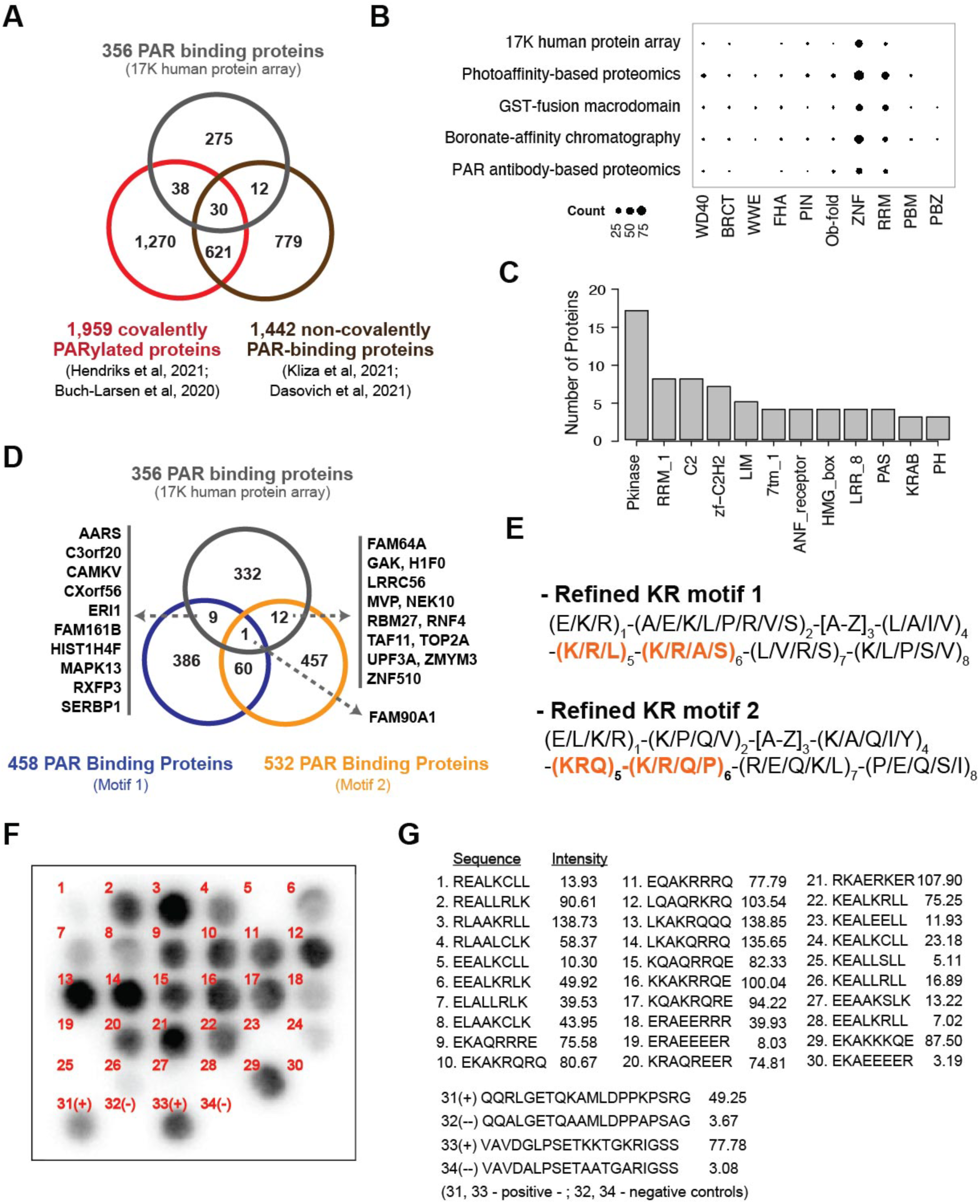
Comparison of PAR-binding protein list with lists from previously published reports and computationally refined KR-based motif. **(A)** A Venn diagram comparing PAR-binding protein lists obtained across four most recent different experimental approaches - MS datasets of (Hendriks *et al*., 2021) and (Buch-Larsen *et al*., 2020) as representative datasets of covalently PARylated proteins, and the datasets of (Kliza *et al*., 2021) and (Dasovich *et al*., 2021) as representative datasets of non- covalent PAR-binding proteins. (B) Dot plot depicting the distribution of domains and motifs identified from our 17K human protein array and four major screening methods – Photoaffinity-based proteomics, GST-fusion macrodomain screen, boronate-affinity chromatography screen, and PAR antibody-based proteomics. (C) A bar graph showing 12 most common Pfam domain families based on the number of occurrences within the 356 PAR-binding proteins. (D) A Venn diagram comparing our experimentally identified protein data lists and computationally predicted PAR-binding protein from previous studies, showing 1 protein (FAM90A1) across the three studies. (E) Two computationally refined versions of previously proposed motifs. Modifications were made to further specify certain regions that were previously unknown, especially the KR-based sequence motifs. (F) A dot blot using different synthetic peptides of KR- based PAR-binding motifs and (G) corresponding PAR-binding intensities including the positive and negative controls of dot blot in spot 31-34.

Comparison of the 12 most common protein domain families (Pfam) from the 356 PAR-binding proteins to computationally predicted data from two prior studies (Gagne *et al*., 2008; Pleschke et al., 2000) identified that kinase domain (Pkinase), RRM_1, C2, zinc finger domain (zf-C2H2), LIM, 7tm_1, ANF_receptor, HMG_box, LRR_8, PAS, KRAB and PH domains were common in our data and each of the theoretical predictions (Figure 2C and Table S7). A Venn diagram of the computationally predicted list of PAR-binding proteins from these prior studies (Gagne *et al*., 2008; Pleschke *et al*., 2000) and our experimental data shows that only 1 protein, FAM90A1, is correctly predicted to be an actual PAR-binding protein across all three methods (Figure 2D and Table S8).

Based on the PAR-binding proteins identified by the protein chip method, we computationally refined each of the two previously proposed KR motifs (Gagne *et al*., 2008; Pleschke *et al*., 2000) to increase specificity as well as diversity (Figures 2E, S2, and Table S9). Motif 1 (Pleschke *et al*., 2000) [AVILMFYW]1-X2-[KR]3-X4 -[AVILMFYW]5- [AVILMFYW]6 -[KR]7-[KR]8-[AVILMFYW]9 -[AVILMFYW]10-[KR]11 was refined to [EKR]1- [AEKLPRVS]2-[A-Z]3-[LAIV]4-[KRL]5-[KRAS]6-[LVRS]7-[KLPSV]8 (Figures 2E and S2A).

Motif 2 (Gagne *et al*., 2008) [HKR]1-X2-X3-[AIQVY]4-[KR]5 -[KR]6-[AILV]7-[FILPV]8 was refined to [ELKR]1-[KPQV]2-[A-Z]3-[KAQIY]4-[KRQ]5-[KRQP]6-[REQKL]7-[PEQSI]8 (Figures 2E and S2B). Overall, these [KR]n[KR]n+1 sequences, which we refer to as KR- based motifs, were commonly identified as part of the computationally predicted motifs.

To examine the PAR-binding capability of the KR-based motifs, a protein spot array with different synthetic peptide combinations based on the two refined motifs (Figures 2F and 2G) was utilized. Motif sequences composed mostly of or dominated by positively charged amino acid R groups – spot 3 (RLAAKRLL), 12 (LQAQRKRQ), 13 (LKAKRQQQ), 14 (LKAKQRRQ), 16 (KKAKRRQE), and 21 (RKAERKER) – showed higher PAR-binding intensities than other spots (Figures 2F and 2G). On the other hand, motif sequences composed mostly of or dominated by negatively charged or neutral amino acid R groups – spot 5 (EEALKCLL), 19 (ERAEEEER), 23 (KEALEELL), 24 (KEALKCLL), 25 (KEALLSLL), 26 (KEALLRLL), 27 (EEAAKSLK), 28 (EEALKRLL), and 30 (EKAEEEER) – showed significantly lower PAR-binding intensities than other spots (Figures 2F and 2G). Specifically, peptide sequences containing glutamic acids consecutively or in high proportion – spot 19 (ERAEEEER) and 30 (EKAEEEER) – corresponded to low PAR-binding densities (Figures 2F and 2G). Two positive (QQRLGETQKAMLDPPKPSRG and VAVDGLPSETKKTGKRIGSS) and negative (QQALGETQAAMLDPPAPSAG and VAVDALPSETAATGARIGSS) controls were applied to normalize the intensity of each spot (Figure 2G). The results taken together suggest that new refined motifs may have greater predictive power as validated in the next section.

### Identification of CPxC and CNxC as PAR-binding motifs

A search for new motifs that may be crucial in PAR-binding using two different sequence alignment methods identified two novel PAR-binding motifs: [C]1-[P]2-x3-[C]4 (CPxC) and [C]1-[N]2-x3-[C]4 (CNxC). The CNxC and CPxC motifs were identified via a modified algorithm of MEME (Figures 3A and S3A). Twelve PAR readers were assigned to the CNxC motif (E-value of 2.4e-003) while 23 other PAR-binding proteins were assigned to the CNxC motif (Figures 3A and S3A). The modified MEGA7 sequence alignment protocol identified 5 common domains in PAR-binding proteins: C2H2-type zinc finger domain (Zf-C2H2), glycine repeat domain, proline-rich domain, glutamine-rich domain, and glutamate-rich domain (Figure 3B and S3B). Among them, Zf-C2H2 had a considerably lower E-value (8.2e-696) than other motifs identified by MEME. High-scoring motifs have a high chance to be functionally linked to the mechanism of protein-PAR recognition. Most of the canonical Zf-C2H2 domain typically contains a repeated 28–30 amino acid sequence that integrates the conserved CNxC and CPxC motifs in their N-terminal part. To validate if each motif contributes to PAR-binding, a peptide spot array with customized peptide combinations containing each motif was performed (Figures S3C and S3D). While the peptide containing the CNxC motif did not exhibit high PAR-binding density, the peptide containing CPxC and KR sequences showed high binding densities (Figures S3C and S3D). The highest binding peptides were spot 3 (TCPTCRK) and spot 4 (TCPTCRKTCPTCRK) (Figure S3D). An increase of PAR-binding intensity was observed with a duplicated CPxC+KR sequence, suggesting a potential constructive interaction between these motifs and PAR (Figures S3C and S3D). Other common motifs that were identified via modified MEGA7 did not exhibit PAR-binding (Figures S3C and S3D).

**Figure 3.**
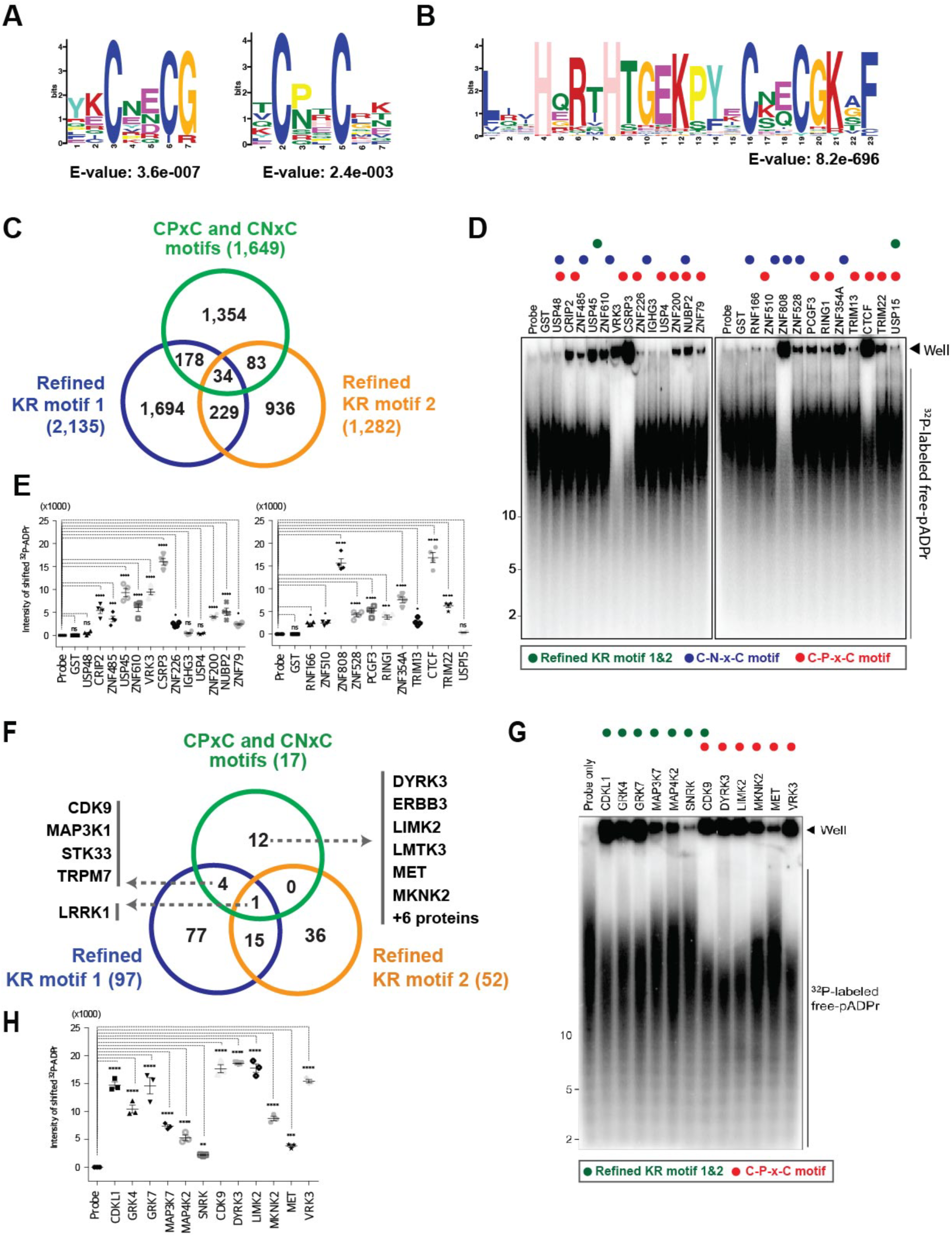
Identification of new PAR-binding Motifs (PBMs), CPxC and CNxC. **(A)** Amino Acid sequence representations for the 2 most significant motifs, CPxC and CNxC, which are novel PAR-binding domains, using MEME *de novo* amino acid alignment protocol. Associated E-values were 2.4e-003 and 3.6e-007 for each CPxC and CNxC motif. (B) Amino Acid sequence representation for the most significant motif observed across PAR-binding proteins using MEGA7 protocol. Associated E-value was 8.2e-696. (C) A Venn diagram comparing human proteins containing new CPxC/CNxC motifs and two refined KR motifs, showing that 34 proteins contain all three motifs. (D) A native electromobility gel shift assay for the validation of PAR-binding of newly predicted proteins based on the new CPxC/CNxC motifs, and (E) a graph of corresponding intensities of the gel shift assay. (F) A Venn diagram comparing human kinases with new motifs and two refined motifs, showing that 1 kinase contain all three motifs. (G) A native electromobility gel shift assay for the validation of PAR-binding of newly predicted kinases based on the new motifs, and (H) a graph of corresponding intensities of the gel shift assay.

Computationally, 1,649 proteins contained the CPxC and/or CNxC motif, 2,135 proteins contained the refined KR motif 1 and 1,282 proteins contained the refined KR motif 2 (Figure 3C and Table S10). 34 proteins were identified to have all three motifs (Figure 3C and Table S10). Being aware that small synthetic peptides, taken out of the protein fold context, might not recapitulate endogenous protein affinities to PAR, electromobility-shift assay with full length proteins was used to validate if CPxC, CNxC, and the refined KR sequence are crucial for PAR-binding (Figure 3D). Randomly selected proteins each containing refined KR, CPxC, and CNxC motifs were purified from *E.coli* and the potential interaction with [^32^P]-labelled PAR was evaluated using a native gel to avoid denaturation of proteins. The assay with quantification of signal confirmed PAR-binding for refined KR-based motif-containing proteins as well as for CPxC and CNXC motif-containing proteins (Figures 3D and 3E). High PAR-binding density was observed in KR-free CPxC and CNxC motif-containing proteins (Figures 3D and 3E).

A group of proteins that contained the CPxC and/or CNxC motif, the refined KR motif 1 and refined KR motif 2 were assigned as kinases (Figure 3F and Table S11).

Seventeen human kinases contained the CPxC and/or CNxC motif, and 97 and 52 kinases contained the refined KR motif 1 and 2, respectively. One kinase, leucine-rich repeat serine/threonine-protein kinase 1 (LRRK1), contained all three motifs (Figure 3F and Table S11). An electromobility-shift assay coupled with quantification of signal using 12 randomly selected kinases confirmed that these kinases bind PAR and contained either refined KR motif 1/2 or CPxC motif (Figure 3G and 3H).

### CPxC and CNxC motifs specify gene ontology of PAR-binding proteins

Characterization of the CPxC and CNxC motifs suggests that they categorize proteins into different families of biological function. 1,007 proteins contained the CPxC motif, and 515 proteins contained the CNxC motif exclusively, and 127 proteins contained both motifs (Figure 4A and Table S12). Gene ontology analysis for proteins containing the CPxC motif indicated that protein ubiquitination was the most prominent function across biological process, molecular function, KEGG pathway categories, whereas for CNxC motif-containing proteins transcription-related functions were most prominent overall (Figure 4B and Table S13). These observations suggest that CPxC-containing proteins are likely to be E3 ubiquitin ligases and CNxC-containing proteins have zinc finger domains for DNA recognition. Representative E3 ligases and zinc finger proteins for CPxC and CNxC motif-containing proteins, respectively are provided in Figure 4C.

**Figure 4.**
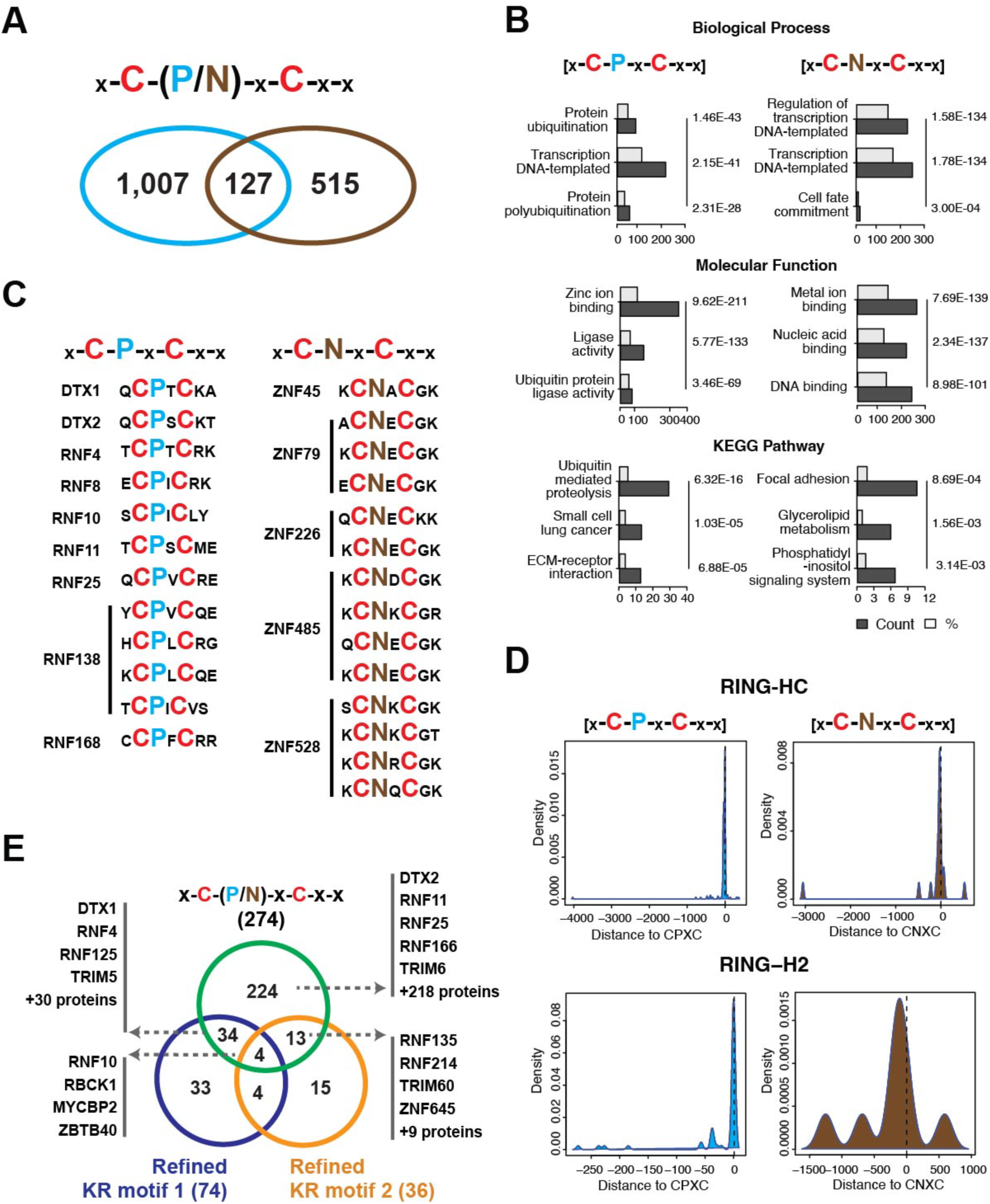
Characterization of the novel PAR-binding Motifs, CPxC and CNxC. **(A)** A Venn diagram comparing human proteins containing the two new PAR-binding motifs, CPxC and CNxC, showing a 127-protein overlap across the two. (B) A bar chart showing the 3 most significant biological processes, molecular functions, and KEGG pathways for both CPxC and CNxC motifs. (C) Two representative protein lists showing some of the proteins found containing the motif CPxC and CNxC, respectively. (D) A distance-based graph of motifs CPxC and CNxC with respect to two protein domains, Zinc finger (RING-HC) and E3 Ligase (RING-H2). (E) A Venn diagram of 614 human protein E3 Ligases containing one or more of PAR-binding motifs: Refined KR motif 1, Refined KR motif 2, and CPxC/CNxC Motif. 4 E3 Ligases contained all three motifs.

Next the location of each CPxC and CNxC motifs in LIM, PHD, RING-H2, and RING-HC domains was examined. The CPxC motif was relatively localized in the same area within the RING domain across different E3 ligases while the location of the CNxC motif tended to vary across different ZNF proteins (Figures S4A and S4B). The distance plot showed that the distances of the RING-HC domain to each of the motifs were proximal to 0, suggesting that each of the motifs are contained within the RING-HC domain (Figure 4D). CPxC was found in many different locations for LIM and PHD domains, but for RING-H2 (E3 Ligase) and RING-HC (Zinc Finger) domain, CPxC was consistently observed in one location (Figures 4D and S4C). On the other hand, the CNxC motif was present in many different locations for LIM, PHD, and RING-H2 domains, but not for the domain RING-HC (Figure S4D). This difference of motif presence in RING-HC suggested that the RING domain as well as the corresponding biological function can be differentiated based on the CPxC or CNxC motif and that CPxC may be a conserved motif for E3 ligases. Along these lines, 384 E3 ligases out of 614 human E3 ligases contained at least one or more of the PAR-binding motifs. 74 and 36 E3 ligases contained the refined KR motif 1 and 2, respectively; 274 E3 ligases contained the specifically CPxC motif; and 4 E3 ligases - RNF10, RBCK1, MYCBP2 and ZBTB40 - contained all three motifs (Figure 4E and Table S14).

### Identification of PAR-binding E3 ligases

Different PAR domain containing E3 ligases (Figure S5A) were selected and analyzed for PAR-binding via a PAR electromobility shift assay (Figure S5B). DTX1, DTX2, RNF4, RNF10, RNF25, HERC3, WWP1 and WWP2 were confirmed to be PAR-binding proteins. RNF11 and RNF12 did not exhibit PAR-binding in this assay (Figure S5B).

DTX1, DTX2, RNF4, RNF10, RNF11, RNF12, RNF25, RNF138, RNF168, HERC3, WWP1 and WWP2 were screened in a double stand break (DSB) reporter system assay (Tang et al., 2013) (Figure S5C). Each E3 ligase was EGFP-tagged and transfected in the DSB reporter cell line the colocalization with the mcherry-LacI-FOKI fusion protein was monitored at the DSB site. Six PAR-binding E3 ligases (DTX1, DTX2, RNF4, RNF8, RNF138 and RNF168) were clearly observed at the DSB site. (Figure S5C).

### CPxC is a PAR-binding motif in PAR-binding E3 ligases, and the RING domain is crucial for PAR-binding E3 ligases

Two groups of E3 ligases were examined. DTX1, DTX2, and RNF4 were chosen to represent group 1 and contained previously identified PAR-binding Motifs (PBM), in addition to the CPxC motif. RNF8, RNF138, and RNF168 were chosen to represent group 2 as they only contained the CPxC motif. DTX1 and DTX2 contain tandem WWE PBM domains similar to RNF146 (Iduna) (DaRosa et al., 2015; Kang et al., 2011). To compare the PAR-binding density between the WWE PBM and the CPXC PBM in the RING domain, we created a double mutant in WWE domain 1 and 2 (Dmut) via site-directed mutagenesis and compared the change in PAR-binding caused by a mutation in the CPxC (CCmut) domain to wildtype via EMSA (Figure 5A). For DTX1, alanine substitutions in 93R/170R were performed to generate Dmut and in 468C/471C for the CCmut. For DTX2, alanine substitutions in 93R/170R were performed to generate Dmut and in 469C/472C for CCmut (Figure 5A). Recombinant wildtype, PBM and CC mutant were incubated with [^32^P]-PAR and separated on a 20% TBE gel. Both [^32^P]-PAR bound signal and unbound free [^32^P]-PAR signal were visualized by autoradiography.

**Figure 5.**
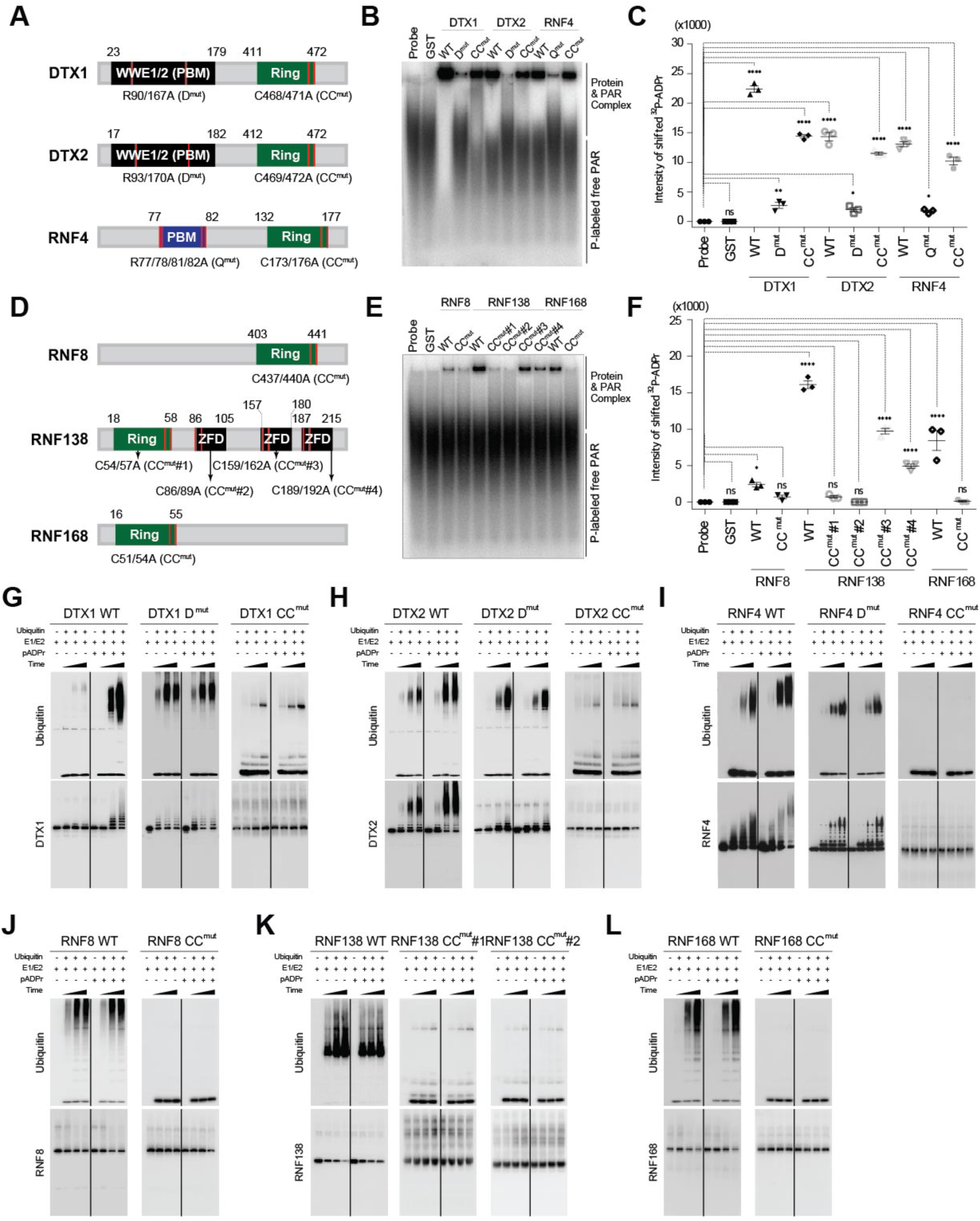
Identification of the PAR-binding motifs in PAR-binding ubiquitin E3 Ligases. (A) Schematic illustrations of PAR-binding motifs in DTX1, DTX2 and RNF4. Each amino acid indicated in red bar were replaced with Alanine by site-directed mutagenesis. (B) Each of recombinant wildtype, PBM and CC mutant was incubated with [^32^P]-PAR and separated in 20% TBE gel. Both [^32^P]-PAR bound signal and unbound free [^32^P]-PAR signal were visualized by autoradiography. (C) Intensity of each [^32^P]-PAR bound protein was quantified and plotted. Gel shift assays were performed in triplicate. \**P*<0.05, \*\**P*<0.01, \*\*\*\**P*<0.0001, by one-way ANOVA when compared between indicated groups by Bonferroni’s posttest, *n.s.* not significant. (D) Schematic illustrations of PAR-binding motifs in RNF8, RNF138 and RNF168. Each amino acid indicated in red bar were replaced with Alanine by site-directed mutagenesis. (E) Both [^32^P]-PAR bound signal and unbound free [^32^P]-PAR signal were visualized by autoradiography. (F) Intensity of each [^32^P]-PAR bound protein was quantified and plotted. Each signal was normalized by GST binding signal. Gel shift assays were performed in triplicate. \**P*<0.05, \*\*\*\**P*<0.0001 by one-way ANOVA when compared between indicated groups by Bonferroni’s posttest, *n.s.* not significant. *n.s.* not significant. (G-F) Each of recombinant wildtype, PBMs and CC mutants of (G) DTX1, (H) DTX2, (I) RNF4, (J) RNF8, (K) RNF138 and (L) RNF168 was subjected to the *in vitro* ubiquitination assay followed by Western blot. Each of recombinant E1, E2, E3, - Ubiquitin and pADPr was added as indicated and incubated as indicated time point.

The Dmut markedly and significantly reduced PAR-binding while the CCmut exhibited a mild, but significant decrease in binding for both DTX1 and DTX2 (Figures 5B and 5C). RNF4 contains a PBM based on the refined KR motif 1. R77, R78, R81 and R82 were selected and substituted into Ala to generate a RNF4 PAR-binding mutant, Qmut (Figure 5A). The Qmut in RNF4 markedly and significantly reduced PAR-binding while the CCmut exhibited a mild, but significant decrease in binding (Figures 5B and 5C).

Alanine substitutions in 437C/440C were performed to generate a PBM mutant, CCmut in RNF8 (Figure 5D). For RNF168, alanine substitutions in 51C/54C were performed to generate a PBM mutant, CCmut. RNF8 and RNF168 CCmuts showed a substantial and significant decrease in PAR-binding, confirming CPxC as a PBM (Figures 5D, 5E, and 5F). RNF138 contained multiple CPxC PBMs in four different locations including the RING domain and the zinc finger domains (ZFD) (Figure 5D). Alanine substitutions in the CPxC motif were made in the RING domain, 54C/57C (CC_#1) or the ZFD domains, 86C/89C (CC_#2) or 159C/162C (CC_#3) or 189C/192C (CC_#4) (Figure 5D). The CC_#1 in the RING domain or the CC_#2 in the first ZFD markedly and significantly reduced PAR-binding while CC_#3 or CC_#4 exhibited a mild, but significant decrease in PAR-binding (Figures 5E and 5F). These results taken together indicate that DTX1, DTX2, RNF4, RNF8, RNF138 and RNF168 are PAR-binding proteins and that the CPxC motif is a PBM domain.

Ubiquitin E3 ligase activities were measured in the presence or absence of PAR polymers in an *in vitro* ubiquitination assay (Figures 5G to 5L). Auto-ubiquitination activity of WT DTX1, DTX2 or RNF4 was increased by addition of PAR polymers in a dose dependent manner (Figures 5G, 5H and 5I). The Dmut in DTX1 or DTX2 increased the autoubiquitination but there was no additional increase in the presence of PAR (Figure 5G and 5H). The CCmut in DTX1 or DTX2 substantially reduces the PAR activation of DTX1 or DTX2 autoubiquitination (Figure 5G and 5H). The Qmut in RNF4 decreased the autoubiquitination and there was no additional increase in the presence of PAR (Figure 5I). The CCmut in RNF4 eliminated autoubiquitination (Figure 6I).

**Figure 6.**
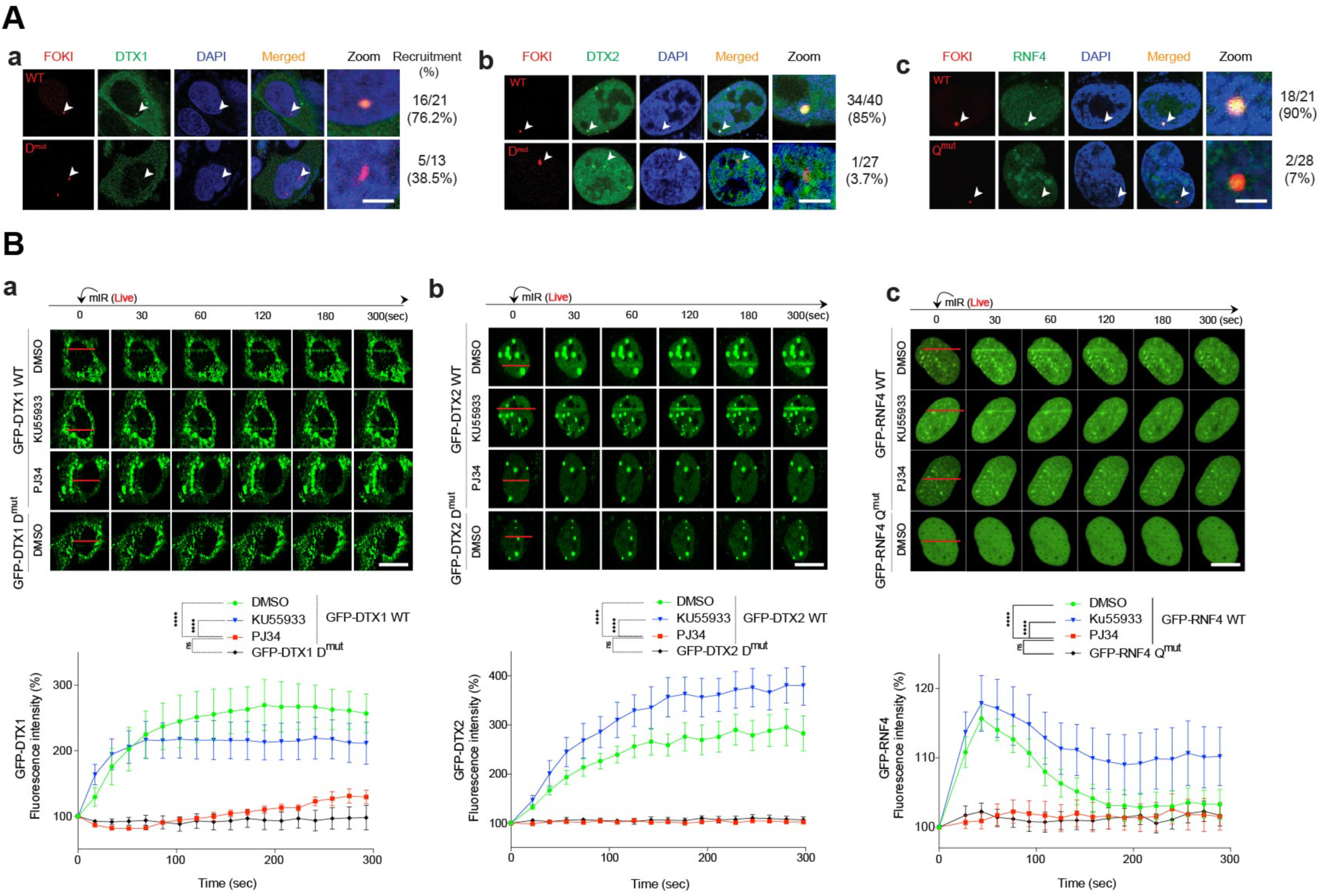
PAR-binding to PAR-binding ubiquitin E3 Ligases is required for the translocation into the damaged region of DNA. (A-C) Identification of co-localization between mCherry-FOKI nuclease and each of GFP-fused wildtype- or PBM of DTX1 (A), DTX2 (B) and RNF4 (C) at DSB site. (D-F) The kinetics of GFP-fused DTX1 (D), DTX2 (E) and RNF4 (B) to DNA lesions in a PARP1 dependent manner. Each of wildtype form of GFP-fused E3s were tested in the presence or absence of PARP inhibitor, PJ34 or ATM inhibitor, KU55933. For PBM mutants for DTX1 and DTX2, gRNA-DTX1 and gRNA-DTX2 was applied before the assay, respectively. Each of PBM form of GFP-fused E3s were tested in the absence of inhibitor. Fluorescence intensity at the laser strips were quantified and plotted as mean ± s.e.m of six independent cells for DTX1 and DTX2 (N=6) or five independent cells for RNF4 (N=5). \*\*\*\**P*<0.0001 by one-way ANOVA when compared between indicated groups by Bonferroni’s posttest. Scale bars, 10 μm.

Auto-ubiquitination activity of WT RNF8, RNF138 or RNF168 was not substantially changed by addition of PAR polymers (Figures 5J, 5K and 5L). The CCmut in RNF8, RNF138 or RNF168 abolished autoubiquitination activity (Figures 5J, 5K and 5L).

These results taken together suggest that DTX1, DTX2 and RNF4 are PAR dependent E3 ligases while the E3 ligase activity of RNF8, RNF138 or RNF168 are not regulated by PAR at least in the assays utilized here.

### Translocation of PAR-binding E3 ligases to DNA damage is PAR-binding and PARP1 dependent

Since the Group 1, PAR-binding E3 ligases, DTX1, DTX2, and RNF4 exhibit PAR dependent E3 ligase activity, WT DTX1, DTX2, and RNF4 and respective PAR-binding mutants were examined in a DSB reporter system assay (Tang *et al*., 2013). Each E3 ligase and their respective PAR-binding mutants were EGFP-tagged and transfected in the DSB reporter cell line and the colocalization with the mcherry-LacI-FOKI fusion protein was monitored at the DSB site. Shown in Figure S5, WT RNF4, DTX1 and DTX2 co-localized at the DSB site, while the PBM mutants for RNF4 (Qmut), DTX1 (Dmut) and DTX2 (Dmut) showed substantially reduced co-localization at the DSB site (Figure 6A). Next, translocation patterns of GFP-tagged WT or PBM mutants for RNF4 (Qmut), DTX1 (Dmut) and DTX2 (Dmut) were examined in HeLa cells after laser microirradiation by fluorescence intensity recording under different conditions over 300 seconds.

Accumulation of GFP-tagged WT RNF4, DTX1 or DTX2 was observed in cells treated with DMSO or the ATM inhibitor, KU55933, but translocation was not detected in cells treated with PARP1 inhibitor, PJ-34 or in cells expressing PBM mutants for RNF4 (Qmut), DTX1 (Dmut) and DTX2 (Dmut) (Figure 6B and 6C). The translocation patterns of GFP-tagged WT or PBM mutants for RNF4 (Qmut), DTX1 (Dmut), DTX2 (Dmut), RNF8 (CCmut), RNF138 (CC_#1mut and CC_#2mut) and RNG168 (CCmut) were also examined in WT HeLa cells and PARP-1 KO HeLa after laser microirradiation by fluorescence intensity recording over 300 seconds (Figure S6A to S6F). Accumulation of GFP-tagged WT DTX1, DTX2, RNF4, RNF8, RNF138 and RNF168 was observed in cells treated with microirradiation while accumulation was eliminated in PARP-1 KO cells (Figure S6A to S6F). PBM mutants of DTX1, DTX2, RNF4, RNF8, RNF138 or RNF168 also failed to translocate (Figure S6A to S6F).

### PAR-dependent E3 Ligases are essential for cell survival and involved in both NHEJ and HR pathways

To examine the role of the PAR-dependent E3 ligases DTX1, DTX2 or RNF4 in cell survival, we performed clonogenic assays to analyze the survival rates of HeLa cells under DTX1, DTX2 or RNF4 knockdown conditions. Doxycycline (DOX) inducible guide RNA resistant constructs (gRNAResDTX1 -WT, -PBM) combined with lentiviral gRNA infection were used to regulate DTX1 and DTX2 expression via the presence (DOX+) or absence (DOX-) of doxycycline (Figure 7Aa-b). DNA damage via ionizing irradiation (IR) was introduced at day 3 and colony formation was monitored over an 18-day period.

Cell survival rates were measured under three different Gy conditions of damage for DTX1 and DTX2. For RNF4, instead of DOX-lenti-gRNA knockdown, HeLa cells were transfected with siRNA-resistant RNF4 constructs and then infected with siRNA (Figure 7Ac). This enabled knockdown of endogenous RNF4, thereby allowing us to focus on the RNF4 variant of interest. For DTX1, we observed that in absence of doxycycline, DTX1 was not expressed, and cell survival decreased in correlation to increasing IR dosage. DTX1 PBM mutant Dmut in presence of DOX exhibited a similar cell survival rate as DOX- condition. On the other hand, WT DTX1 in presence of DOX showed higher cell survival rate (Figure 7Aa). Similar results were observed for DTX2, where WT DTX2 DOX-, DTX2 Dmut DOX-, and DTX2 Dmut DOX+ showed a lower cell survival rate than DTX-WT DOX+ (Figure 7Ab). For RNF4, V5 non-siRNA-resistant RNF4 knocked down with siRNF4, and siRNA-resistant-Qmut showed decreased cell survival rates. siRNA resistant RNF4-WT showed higher cell survival rate (Figure 7Ac).

**Figure 7.**
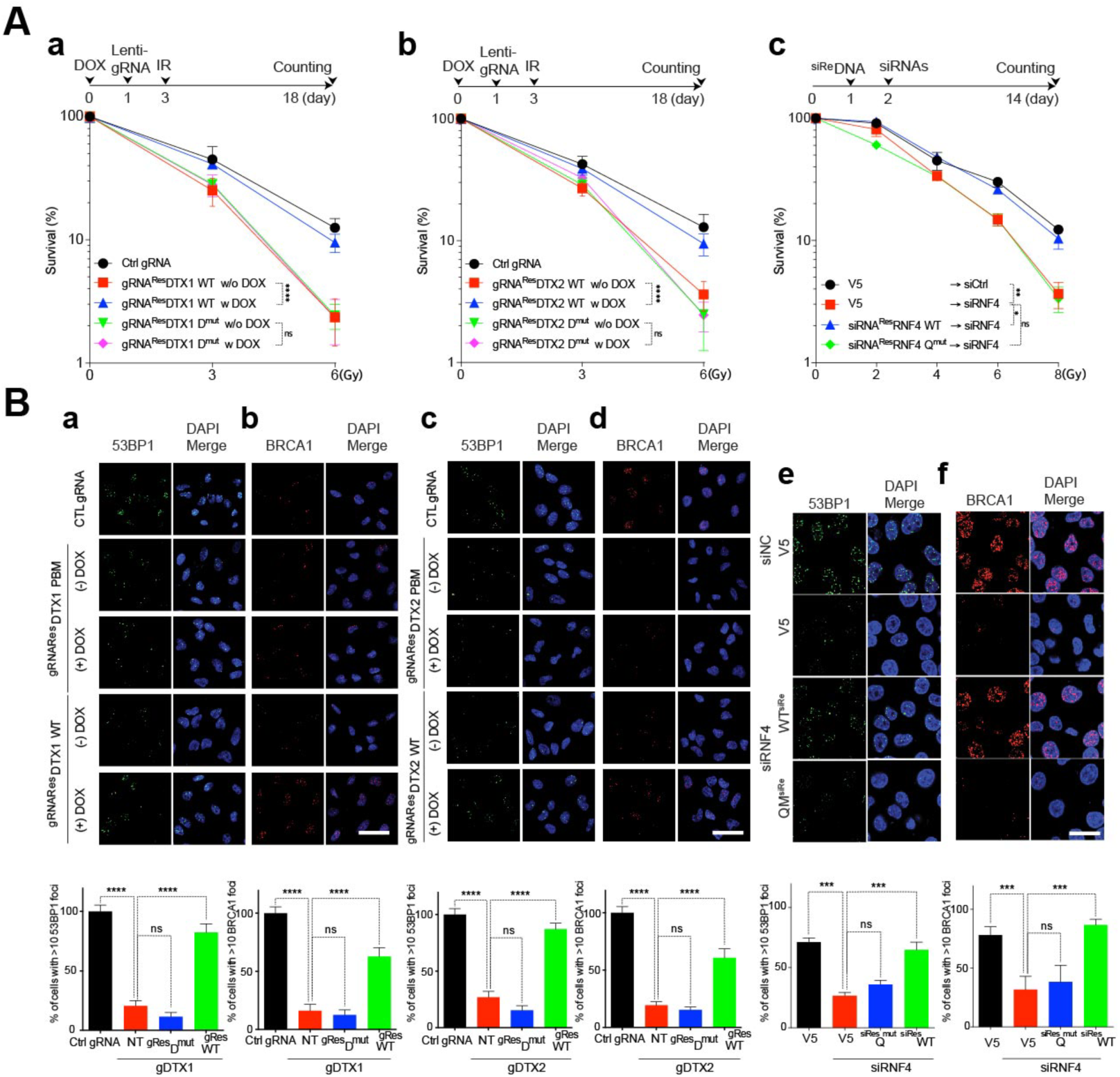
PAR-binding E3 Ligases are essential for cell survival and involved in both NHEJ and HR pathway. (A) Clonogenic analysis. (Aa) HeLa cells were transfected with doxycycline-inducible gRNA resistant constructs (gRNAResDTX1 WT or gRNAResDTX1 Dmut) and induced in the presence or absence of doxycycline and then, cells were infected with Lentiviral gRNA for DTX1. Followed by ionizing irradiation with different dosages, the colony formation assay was performed as indicated. (Ab) HeLa cells were transfected with doxycycline-inducible gRNA resistant constructs (gRNAResDTX2 WT or gRNAResDTX2 Dmut) and induced in the presence or absence of doxycycline and then, cells were infected with Lentiviral gRNA for DTX2. Followed by ionizing irradiation with different dosages, the colony formation assay was performed as indicated. (Ac) HeLa cells were transfected with siRNA resistant RNF4 constructs (siResRNF4 WT or siResRNF4 Qmut) and then transfected with siRNA. Followed by ionizing irradiation with different dosages, the colony formation assay was performed as indicated. \**P*<0.05, \*\**P*<0.01, \*\*\*\**P*<0.0001, by one-way ANOVA when compared between indicated groups by Bonferroni’s posttest, *n.s.* not significant. (B) Representative images or frequency of foci formation of 53BP1 (Ba, Bc, Be) and BRCA1 (Bb, Bd, Bf) at DNA break sites after irradiation. Depletion of endogenous E3s and ectopic expression gRNA- or siRNA-resistant wildtype or PBM form of E3s were prepared as indicated. Followed by immunocytochemistry, numbers of cells with at least 10 foci were counted and plotted. \*\*\**P*<0.001, \*\*\*\**P*<0.0001, by one-way ANOVA when compared between indicated groups by Bonferroni’s posttest, *n.s.* not significant. Scale bars, 10 μm.

To further investigate the roles of these PAR-dependent E3 ligases in specific DDR pathways, HR and NHEJ reporter systems were employed (Figure S7A) (Gunn and Stark, 2012). DTX1 knockdown cells showed altered HR and NHEJ activities, which were rescued by siRNA-resistant WT DTX1, while siRNA resistant DTX1 Dmut exhibited decreased HR and NHEJ efficiency (Figure S7Ba and S7Bd). DTX2 knockdown cells also showed altered HR and NHEJ activities that were rescued by siRNA resistant WT DTX2; both HR and NHEJ efficiency of DTX2 Dmut decreased slightly (Figure S7Bb and S7Be). RNF4-knockdown cells showed similar effects and were rescued by reconstitution of siRNA-resistant WT RNF4, but not or RNF4 Qmut (Figure S7Bc and S7Bf). Next IR-induced foci formation assays for both 53BP1 and BRCA1 were performed. Knockdown of DTX1, DTX2 or RNF4 inhibited both IR-induced 53BP1 (Figure 7Ba, 7Bc, 7Be) and BRCA1 foci formation (Figure 7Bb, 7Bd, 7Bf) that was rescued by reconstitution of WT but not by the PBM mutants. These results indicate that PAR-binding ubiquitin E3 ligases are upstream factors for IR-induced 53BP1 and BRCA1 foci formation and critical for promoting both NHEJ and HR pathways.

## Discussion

In this study, we systematically profiled PAR-protein interactions using a protein microarray-based approach. The PAR-binding proteins that we identified were primarily located in the nucleus, as we would expect for a chromatin regulatory molecule, but a significant number of PAR readers were found to reside in the cytoplasm and in transmembrane regions of the cell. Consistent with the known role of PAR signaling in DNA damage repair and transcriptional regulation (Alemasova and Lavrik, 2019; Liu *et al*., 2017; Malanga et al., 1998; Pleschke *et al*., 2000; Stilmann et al., 2009), we observed that the two most common protein functions related to PAR readers was linked to nucleic acid binding and transcription factor activity.

In line with previously published reports, 80 proteins identified as PAR readers via the protein microarray were previously found in non-covalent PAR binding or covalent ADP-ribosylation datasets (Figure 2A), highlighting the quality of the PAR reader dataset using the protein microarray-based approach. Known PAR-binding domains such as, C2H2-type zinc finger, RRM, or OB-fold domains were predominant modules as PAR readers (Figure 2B). A major advantage of protein microarrays is that they are highly sensitive to targets that might be expressed at a very-low level in cells. In addition, the interactions observed with PAR are direct. This contrasts with cell-based assays where screening for PAR-binding is complexified by protein-protein or protein-DNA/RNA interactions leading to a confusing image, in which it is hard to distinguish non-covalent PAR-binding from covalent PARylation. Thus, the use of protein microarrays allows a robust resolution of this cell-based complexity.

The PBMs were the first motifs found to mediate interaction with PAR (Gagne *et al*., 2008; Pleschke *et al*., 2000). They feature a loose consensus sequence composed of basic amino acids flanked by aliphatic residues. To reach higher selectivity, we refined these motifs and proved that they are highly effective for selecting direct PAR-binding proteins with motif specificity and accuracy as shown in Figure 2F, 2G, 3G. These computationally refined KR-based motifs 1 and 2 also explain the charge-based interaction with PAR. The result of the protein spot array showed that peptides with positively charged sequences bound strongly to PAR, whereas negatively charged peptides didn’t bind to a great extent to PAR (Figure 2F and 2G), which is consistent with PAR being a negatively charged polymer.

Five common modules in PAR-binding proteins were identified including the Zf-C2H2 domain, glycine repeat domain, proline-rich domain, glutamine-rich domain, and glutamate-rich domain via modified MEGA7 sequence protocol (Figure S3B). Zf-C2H2 had a considerably lower E-value (8.2e-696) than other motifs identified by MEME. These domains exhibited minimal PAR-binding on a peptide array suggesting that these may function as PAR readers that is contextually dependent on the architecture of the protein or the protein domain structure, as it is the case for zinc fingers. Two new PBMs, CPxC and CNxC motifs, were identified and validated as PAR readers. The PAR-binding intensity of CNxC appeared to be weaker than CPxC and both increased proportionally to their duplicity, potentially suggesting a constructive interaction between each motif (Figure S3C and S3D). The strong PAR-binding intensity of CPxC or CNxC motif-containing proteins under native conditions suggest that the PAR-binding capability of these motifs may depend on protein structure. Consistent with the notion that CPxC and CNxC PBMs may be structural motifs are the findings that mutations into these motifs decrease PAR-binding intensities on native EMSA but not protein spot arrays. CPxC and CNxC are distinct in terms of their associated proteome.

Interestingly, CPxC and CNxC functionally specify the category of PAR-binding protein family as proteins containing CPxC are likely to be E3 Ubiquitin ligases, and proteins containing CNxC are likely to be zinc finger proteins that serve transcriptional roles.

CPxC was primarily present in RING domains at a consistent location, whereas CNxC was present in zinc fingers with some variations in its position. These observations further suggest that CPxC may be a conserved motif for E3 Ubiquitin ligases and may contribute to PAR-dependent regulation of E3 ligase activities, especially since many E3 ligases contained CPxC or KR-based motifs and showed PAR-binding (Figure 4E). Due to complexity of ZNF and RING containing proteins, these CPxC and CNxC motifs as PAR readers needs to be further explored.

Utilizing these PBM, we were able to identify several new classes or expand classes of proteins that are potentially regulated by PAR-binding. New classes of proteins that should be investigated for regulation by PAR include signaling molecules, transcription factors, nucleic acid binding proteins, calcium binding proteins, ligases, oxidoreductases, enzymes, transferases, hydrolases, and receptors. One particular group were kinases, as some of them contained the CNxC and/or CPxC motifs, the refined KR motif 1 or 2. In this new class of PAR-binding kinases, 17 human kinases contained the CPxC and/or CNxC motif, and 93 and 51 kinases contained the refined KR motif 1 and 2, respectively, with one kinase, leucine-rich repeat serine/threonine-protein kinase 1 (LRRK1), containing all three motifs. We were able to confirm that 12 randomly selected kinases bind PAR, and this may suggest the presence of PAR dependent kinases. Considering the ubiquitous role of kinases in intracellular signaling, studying PAR-dependent kinases may be a point of interest for future studies to understand how PAR-binding to these kinases can regulate their activity, function and/or localization to biological processes. Since kinases contain an ATP binding pocket that can recognize ADP (Becher et al., 2013), it will be important to determine if PAR-binding can regulate kinase function that is separate from the ATP binding pocket.

Ubiquitin E3 ligases represent a class of expanded proteins that bind PAR. 384 E3 ligases out of 614 human E3 ligases contained at least one or more of the PAR-binding motifs. 74 and 36 E3 ligases contained the refined KR motif 1 and 2, respectively, 274 E3 ligases contained the specifically CPxC motif, 4 E3 ligases, RNF10, RBCK1, MYCBP2 and ZBTB40, contained all three motifs. DTX1, DTX2, RNF4, RNF10, RNF25, HERC3, WWP1 and WWP2 were confirmed to be PAR-binding E3 ligases and DTX1, DTX2, RNF4, RNF8, RNF138 and RNF168 were found to translocate to DNA damage sites is PARP-1 and PAR-binding dependent. DTX1, DTX2, RNF4 ubiquitin E3 ligase activity is increased in the presence PAR, and we show that each E3 ligase plays a role in the DDR. RNF4 is a KR-based PBM protein while DTX1 and DTX2 PAR-binding is similar to the WWE PBM in RNF146 (Iduna) (DaRosa *et al*., 2015; Kang *et al*., 2011). All three proteins contained the CPxC motif. We observed a difference in PAR-binding intensity and PAR-dependent E3 ligase contribution between the KR motif or WWE motif and the CPxC motif suggesting that certain proteins have multiple PBMs and can maintain a diversity of function despite interfering with binding on one motif. A topic for future study will be sorting out the role of each motif in the biologic function of proteins containing multiple PBMs. On the other hand, RNF8, RNF138, and RNF168, which only contain the CPxC motif, CPxC was shown to be critical in enabling E3 ligase function. Future studies are required to sort out the importance of this motif in regulating the function of these E3 ligases.

In summary, we report the identification and characterization of new set of PAR-binding proteins and novel PBMs. Our PARylome greatly expand the spectrum of PAR-binding by a variety of new PAR readers and provides striking new insights into the cellular pathways regulated by PAR.

## Acknowledgements

**Funding:** This work was supported by the Daniel Nathans Award to V.L.D. and the Leonard and Madlyn Abramson Professor in Neurodegenerative Diseases to T.M.D. and the National Research Foundation of Korea grants funded by the Korean government (MSIP) [NRF-2020R1A2C2004988] to H.C.K.

## Authors contributions

Conceptualization, B.G.K., S.-U.K., H.C.K.,T.M.D., V.L.D.; Methodology, B.G.K., S.-U.K., H.C.K.; Validation, B.G.K., S.-U.K., H.C.K.; Formal Analysis, B.G.K., S.-U.K., H.C.K..; Investigation, B.G.K., S.-U.K., H.C.K, J.J.K., J.K., S.Y.L., S.K., K.L.S., S.H., J.-P.G.Y.-I.L.; Resources, H.Z., T.M.D., V.L.D.; Writing-Original Draft, B.G.K., S.-U.K.,T.M.D., V.L.D.; Visualization, B.G.K., S.-U.K., H.C.K.,; Supervision, G.G.P., T.M.D., V.L.D.; Funding Acquisition, T.M.D., V.L.D.

## Declaration of interests

The authors declare no competing interests.

## STAR METHODS

### RESOURCE AVAILABILITY

#### Lead Contact

Further information and requests for resources should be directed to and will be fulfilled by the Lead Contact, Ted M. Dawson, M.D., Ph.D.

#### Materials Availability

Further information and requests for resources and reagents listed in Key Resources Table should be directed to the Lead Contact. All unique/stable reagents generated in this study are available from the Lead Contact with a completed Materials Transfer Agreement.

#### Data and Material Availability

All biological resources, antibodies, cell lines and model organisms and tools are either available through commercial sources or the corresponding authors. The data sets generated during and/or analyzed during the current study are available from the corresponding author on reasonable request.

### EXPERIMENTAL MODEL AND SUBJECT DETAILS

#### Materials and Methods

##### Cell lines

HeLa CCL-2 cell line and HEK293FT cell line were purchased from ATCC and Invitrogen, respectively and maintained in DMEM with 10% (v/v) fetal bovine serum (FBS; GIBCO) and 1% Pen/Strep (GIBCO). U2OS-based DR-GFP and EJ5-GFP cells were kindly provided by Dr. Jeremy Stark and maintained with DMEM containing 10% FBS, 1 μg/ml puromycin (GIBCO) and 1% Pen/Strep. U2OS-2-6-3 reporter cells integrated to Lac operator repeats (X256) was kindly provided by Dr. Roger Greenberg and maintained with DMEM containing 10% FBS, 1 μg/ml puromycin, 2 mM L-glutamine and 1% Pen/Strep.

##### Plasmids, site-directed mutagenesis, siRNA, and antibodies

pENTRY Donor 211 vectors of human ubiquitin E3 ligases were purchased from the HIT core facility (Johns Hopkins University) or Harvard ORF collection (Harvard University) or Addgene. For GFP-tagged E3 ligase expression, pDEST53 (GFP N-terminal) vector was purchase and used (Invitrogen). For Dox-inducible expression, pLIX402 destination vector (Addgene #41394) was used. For recombinant protein expression, pDEST15 (Invitrogen, GST N-terminal) or pDEST527 (Addgene, #11518, 6XHis N-terminal), were used respectively. To generate PAR-binding mutant, site-directed mutagenesis was performed using a QuikChange site-directed mutagenesis kit II (Stratagene). siRNA was transfected into cells using Lipofectamine RNAiMAX (Invitrogen). Antibodies, siRNA, and primers used in this study were described in Table S14, respectively.

##### Purification of recombinant PARP1

Human recombinant PARP1 was purified as previously described (Langelier et al., 2008) using expression vectors for his-tagged recombinant human PARP1 (a gift from John M. Pascal). His-tagged PARP1 was expressed in BL21 Gold (DE3) cells (Agilent) in LB medium. Cells were lysed in 25 mM HEPES (pH8.0), 500 mM NaCl, 500 mM TCEP and complete protease inhibitor cocktail (Roche) with high pressure microfluidizer. The cleared cell lysate was loaded into Ni-NTA column (Roche), washed, and eluted in 20 mM NaPO4 (pH7.4), 500 mM NaCl, 500 mM TECEP, 500 mM imidazole (pH7.4) and 1% glycerol. The proteins were then subjected to size exclusion chromatography on a Superdex 200 16/60 column (GE Healthcare) equilibrated in 25 mM HEPES (pH8.0), 0.1 mM TECEP, 150 mM NaCl and the fractions were analyzed by SDS-PAGE gel, pooled, and stored at -80^°^C.

##### Synthesis of biotin and [^32^P]-labeled poly(ADP-ribose) polymer (PAR)

3 ug of recombinant PARP1 was incubated in reaction buffer containing 100 mM Tris-cl, pH 8.0, 10 mM MgCl2, 8 mM DTT, 10% glycerol, 23 ug calf thymus activated DNA, 4 mM biotin-labeled NAD or 75uCi [^32^P]-labeled NAD for 30 min at 30°C. To collect automodified PARP1, 3 M CH3COONa and isopropanol were added in sample thereafter automodified PARP1 was precipitated by centrifugation at 10,000 g for 10 min at 4 ^°^C. Biotin-labeled or [^32^P] labeled PAR was alkali extracted from the automodified PARP-1 and affinity purified using a dihydroxyboronyl BioRex (DHBB) resin. (Menard and Poirier, 1987)

##### Protein microarray

Human 17K protein chips were purchased from CDI Laboratories. Biotinylated RNA positive oligonucleotides (poly A) were purchased from Thermo (Q152P0100). Each protein chip was blocked with 5% skim milk (BD Bioscience) in PBS-T (0.05% Tween 20). After blocking, each protein chip was incubated for 2 hr at room temperature with Biotin-Poly A tail or Biotin-PAR polymer and washed for 3 times in PBS-T (0.05% Tween 20). Anti-GST (CDI Laboratories) or anti-Streptavidin (Sigma) or anti-PAR (Trevigen) were probed at 4°C for 1 hr and washed for 3 times in PBS-T (0.05% Tween 20). Anti-mouse Alexa488 and Anti-rabbit Alexa594 were probed at 4°C for 1 hr and washed for 3 times in PBS-T (0.05% Tween 20). After removal of residual washing buffer by brief centrifugation, each chip was air-dried and scanned with GenePix 4300A (Molecular Devices). For PARG treatment, recombinant human PARG protein (Sigma) was incubated in reaction buffer containing 100 mM Tris-cl, pH 8.0, 15 mM MgCl2, 8 mM DTT, 10% glycerol for 1 hr at 30°C.

##### PAR overlay assay

Protein was electrophoresed by SDS-PAGE and transferred onto NC or PVDF membranes. Each membrane was blocked with 5% skim milk (BD Bioscience) in PBS- T (0.05% Tween 20). After blocking, the membrane was incubated for 2 hr at room temperature with PAR polymer and PAR-binding proteins were detected by anti-PAR antibody. Immunoblots were visualized in X-ray films (AGFA) or ImageQuant LAS 4000 by an ECL method (Thermo).

##### Protein spot array

Customized peptide arrays used in this study were purchased from JPT Peptide Technologies (GmbH, Germany). Briefly, each peptide spots membrane was rinsed with methanol for 5 min to avoid the precipitation of hydrophobic peptides during the following TBS-T washing. Each membrane was blocked with 5% skim milk (BD Bioscience) in TBS-T (0.05% Tween 20). After blocking, the membrane was incubated for 30 mins at room temperature with [^32^P]-PAR polymer and washed thoroughly. Each membrane was exposed to Imaging plate (Fuji film, BAS-MS) and signal was visualized by phosphorimager (GE, Typhoon FLA-9500).

##### Electromobility gel shift assay

Each protein was incubated with purified [^32^P]-labeled PAR in binding buffer (50mM Tris pH7.5, 150 mM NaCl) at RT for 10 mins. Each sample was mixed with DNA loading dye (Thermo) and electrophoresed in 4-20% TBE PAGE (Invitrogen). Each gel was exposed to Imaging plate (Fuji film, BAS-MS) and signal was visualized by phosphorimager (GE, Typhoon FLA-9500).

##### *In vitro* ubiquitination assay

Recombinant E1, UbcH5c and ubiquitin were purchased from Boston Biochem. E1 (50 nM), E2 (50 nM) and E3 protein were incubated with recombinant his-ubiquitin (200 mM) in reaction buffer containing 50 mM Tris-Cl, pH7.5, 2.5 mM MgCl2, 2 mM DTT, 2 mM ATP at 37°C. Reaction was stopped by addition of Laemmli buffer containing beta-mercaptoethanol and subjected into following Western blot. Either His (Thermo) or Pan Ubiquitin antibody (DAKO) was used to detect ubiquitinated protein and free ubiquitin together.

##### Generation of Knock-out (KO) HeLa cell lines

CRISPOR (http://crispor.tefor.net/gRNAs) was used for designing gRNAs and selected gRNAs were subcloned into the LentiCRISPR v2 (Addgene #52961). gRNA expression plasmids were transfected into HeLa cells using jetPrime transfection reagent (Polyplus transfection). After 2 days of transient drug selection by 1ug/ml Puromycin, single cell propagation was carried out by limiting dilution. Expression levels of target proteins in each single cell originated clones were confirmed by Western blot. Empty LentiCRISPR v2 was used for generation of control cell line in the same setting of experimental conditions.

##### Lentiviral preparations for overexpression

Invitrogen ViraPower lentiviral packaging system was employed for high-titer viral preparations for effective transduction. Lentiviral vectors were transfected into HEK 293FT cells along with viral packaging plasmids using calcium phosphate method (Elegheert et al., 2018). After 12 hr, cells were shocked with 10% DMSO in PBS for 2 minutes thereafter cells were further incubated during 18 hr with fresh medium. Viral particles were precipitated by centrifugation at 25,000 g for 3 hr. Pellets were dissolved with serum free medium and stored at -80°C.

##### FOKI assay

Each EGFP-tagged E3 ligases were transfected into DSB reporter cells by using Lipofectamine 3000 at 24 hr before live imaging. To induce the DSB, 4-OHT and Shiled- 1 were pretreated in between 30 mins to 6 hr before observation. (Tang *et al*., 2013).

Colocalization signals was analyzed and visualized by a Zeiss LSM 710 confocal microscope.

##### HR, NHEJ assay

HR or NHEJ efficiency was measured by U2OS-DR-GFP (HR) or U2OS-EJ5-GFP (NHEJ) reporter cell lines, respectively. U2OS-DR-GFP and U2OS-EJ5-GFP cells were transfected with each siRNA. The following day, I-SceI and siRNA-resistant constructs were delivered to each reporter cell, and 72 hr later, they were assayed for GFP-positive by the flow cytometry (BD).

##### UV laser induced DNA damage

For induction of localized DNA damage, HeLa cells were plated onto 25 mm glass bottom culture dishes and each GFP-tagged clone was transfected using jetPrime transfection reagent at 24 - 30 hr before microirradiation and gRNA expression vector was transfected 48 hr before microirradiation, if required. The PARP1 inhibitors PJ34 (5 μM; Santa Cruz Biotechnology) and ATM inhibitor KU55933 (10 μM; Calbiochem) were treated for 1 hr before laser induced DSBs and same concentration of DMSO was treated as a control. Cells were sensitized with 2 µM of Hoechst 33342 Solution (Thermo Fisher Scientific) for 5 min and mounted on a preheated stage at 37°C on a Zeiss LSM 710 confocal microscope equipped with 405-nm laser source in the setting of 405 nm wavelength in 1 literation at 50% output or 30 literation at 100% output at 37°C chamber supplying 5% CO2. The laser was focused on a small rectangular strip of nucleus through 63X oil objective to induce localized DNA damage.

##### Clonogenic survival assay

Clonogenic viability was examined using a colony forming assay. For DTX1 and DTX2, cells were transfected with doxycycline-inducible gRNA resistant WT or PBM constructs and induced in the presence or absence of 2uM of doxycycline for 24 hr. Then, cells were infected with Lentiviral gRNA. At 48 hr after gRNA delivery, cells were irradiated with indicated amount of ionizing radiation dose and further grown for 18 days until growing colonies were visible. For RNF4, cells were transfected with each of siRNA-resistant DNA and maintained for 24 hr. Cells were harvested and plated onto 6-cm dish followed by siRNA transfection and further maintained for 14 days. Resulting colonies were fixed with methanol and stained with 0.5% Crystal violet (Sigma). Colonies were counted and normalized to plating efficiencies.

##### Measurement of 53BP1 and BRCA1 foci formation by immunocytochemistry

For DTX1 and DTX2, cells were transfected with doxycycline-inducible gRNA resistant WT or PBM constructs and induced in the presence or absence of 2uM of doxycycline for 24 hr. Then, each gRNA expressing lentivirus was introduced. At 48 hr after gRNA delivery, cells were irradiated with appropriated amount of ionizing radiation. For RNF4, cells were transfected with each of siRNA-resistant DNA and maintained for 24 hr. Cells were plated onto 6-cm dish followed by siRNA transfection and irradiated with appropriated amount of ionizing radiation. PFA fixed cells were washed and permeablized with PBS containing 0.5% Triton X-100 for 5 min at RT and incubated with blocking buffer (PBS containing 10% Normal Goat Serum, 0.1% Triton X-100) for 1hr at RT. Then incubated with each 3 ug of 53BP1 (Abcam) or BRCA1 (SantaCruz) antibody diluted in Ab buffer (PBS containing 1% Normal Goat Serum, 0.1% Triton X- 100) at 4°C for overnight. The cells were incubated with the indicated secondary antibodies diluted in the PBS containing 0.01% Triton X-100 at room temperature for 1hr at RT. After washing, DAPI was used for nuclear staining and cover slips were mounted to microscope glass slides using fluorescence immunomount media (Fisher Scientific) and visualized on a LSM710 (Carl Zeiss) or A1 (Nikon) confocal microscope. Cells having over 10 foci per nucleus were manually selected and counted.

##### Statistical analysis

Graphs were created, and statistics were calculated using Prism software version 8.4.3 (GraphPad) and noted in the text and figure legends. Data are represented as mean ± s.e.m. Information on sample size (n, given as a number) for each experimental group/condition, statistical methods and measures are available in all relevant figure legends.

##### Computational analysis

**Identification of new motif analysis:** 1,134 and 642 proteins have PAR-binding motifs based on each x-C-P-x-C-x-x and x-C-N-x-C-x-x amino acid sequences were found on 19,728 all protein sequences using in-house made script. X means that the identity of an amino acid is undetermined. The 290 and 60 corresponding proteins were identified by comparing the 1,134 and 642 proteins with 614 human ubiquitin ligases. Then, the distances between two PAR-binding motifs and several protein domains were measured to check if the motifs are located on the domains. **Refine motifs:** Our more stringent PAR-binding motifs than those of previously published paper (Gagne *et al*., 2008) were narrowed the range of amino acids sequences corresponding to each position and were slightly modified. When determining the expected range of amino acids in a position, we used amino acids belonging to the same amino acid group as nonpolar or positively charged group. As a result, 12 PAR-binding motifs with statistical significance of the difference in number of motifs for 356 identified proteins compared to all proteins were computationally identified using in-house made script. The motif 1, 2, 9 among them have been experimentally validated that (experimental explanation).

**Characterization of 356 unique PAR binding proteins using 17k human protein microarray:** The identification of BioGRID (Oughtred et al., 2021) hub proteins and UniProt (UniProt, 2019) sequence features was performed using the Database for Annotation, Visualization and Integrated Discovery (DAVID, version 6.8) (Sherman et al., 2022).

## Supplementary Information

### Supplementary Figures and Figure Legends

**Figure S1.**
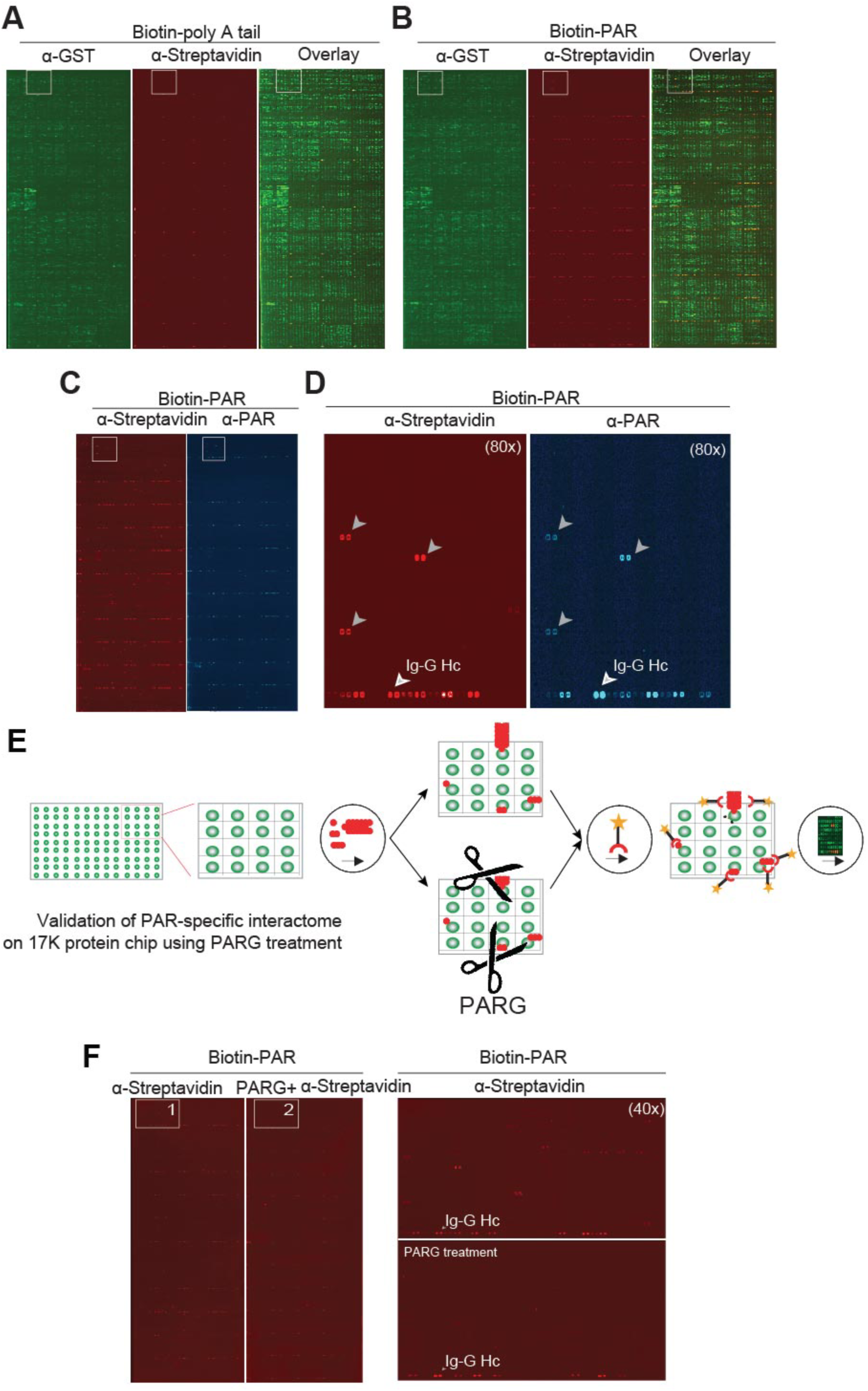
(A) A magnified image from biotin-poly A tail, (B) biotin-PAR, and (C) anti-PAR based detection of PAR-binding proteins as well as (D) its 80x magnified image. (E) Validation of PAR specific interactome on 17K protein chip using PARG treatment and (F) detections of PAR-binding proteins with the addition of anti-streptavidin antibodies.

**Figure S2.**
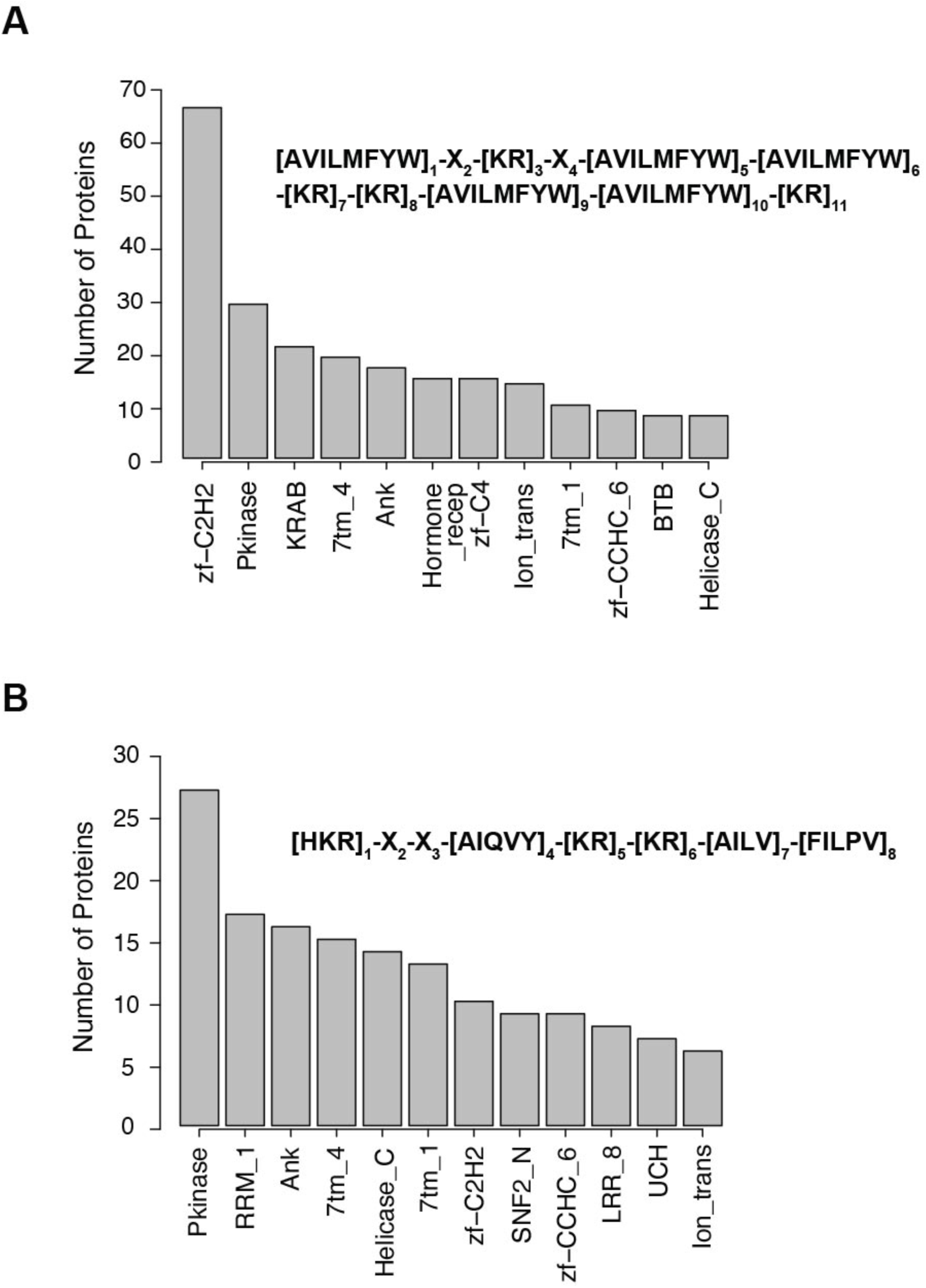
A bar graph showing 12 most common Pfam domain families based on computationally predicted data from (A) (Pleschke *et al*., 2000) and (B) (Gagne *et al*., 2008).

**Figure S3.**
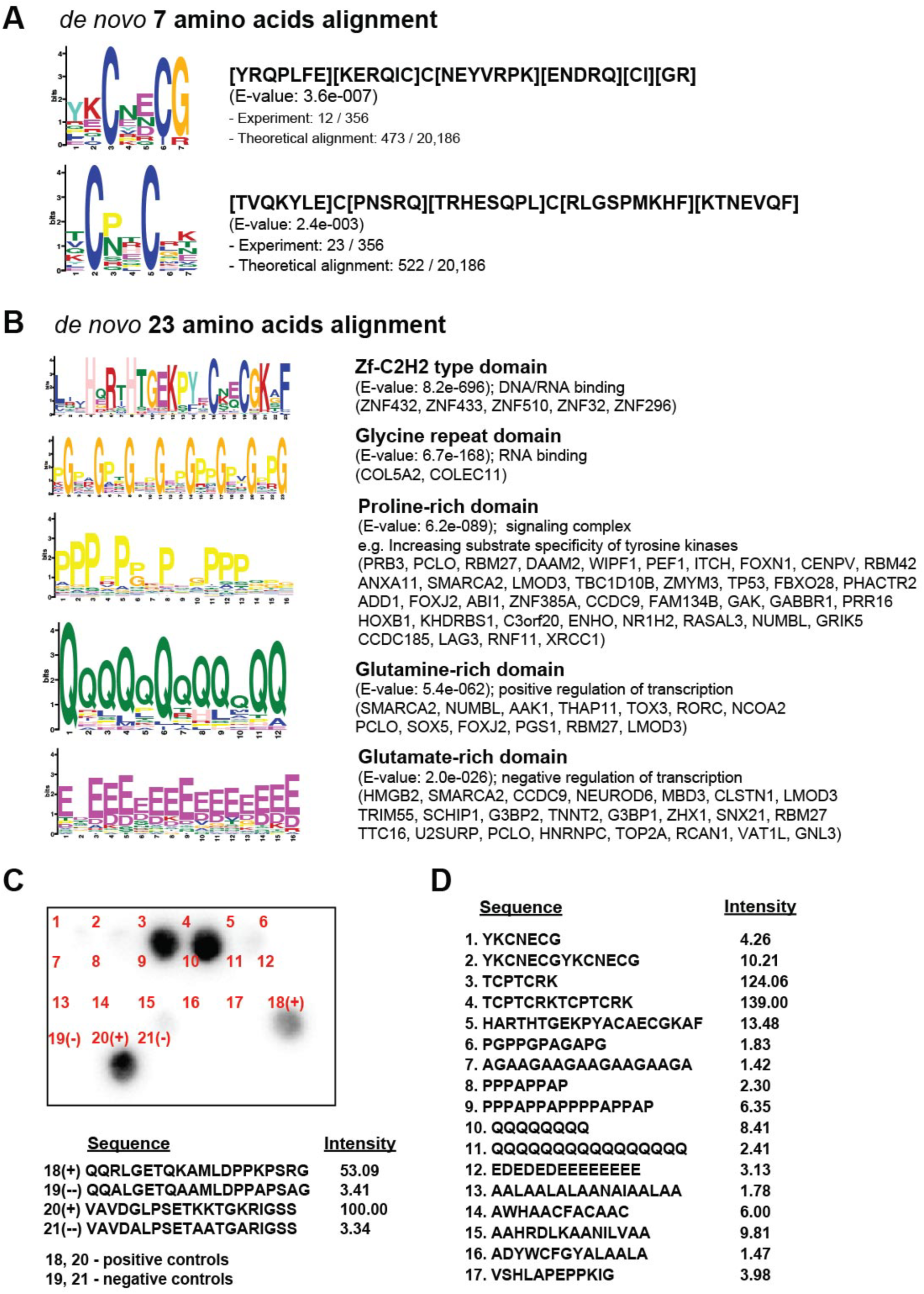
(A) Sequence representations for 2 significant motifs as novel PAR-binding domains using MEME *de novo* amino acid alignment protocol. The experimental data collected predicts the presence of the first sequence in 473 of the 20,186 sequences, while it predicts the presence of the second sequence in 522 of the 20,186. (B) Sequence representations for 5 most significant motifs observed across PAR-binding proteins using MEGA7 protocol. Data such as E-value and domain type function were also evaluated across the 5 domain types. (C) A protein spot array and (D) a table of sequences corresponding to the numbers labeled on the protein spot array and their corresponding intensities.

**Figure S4.**
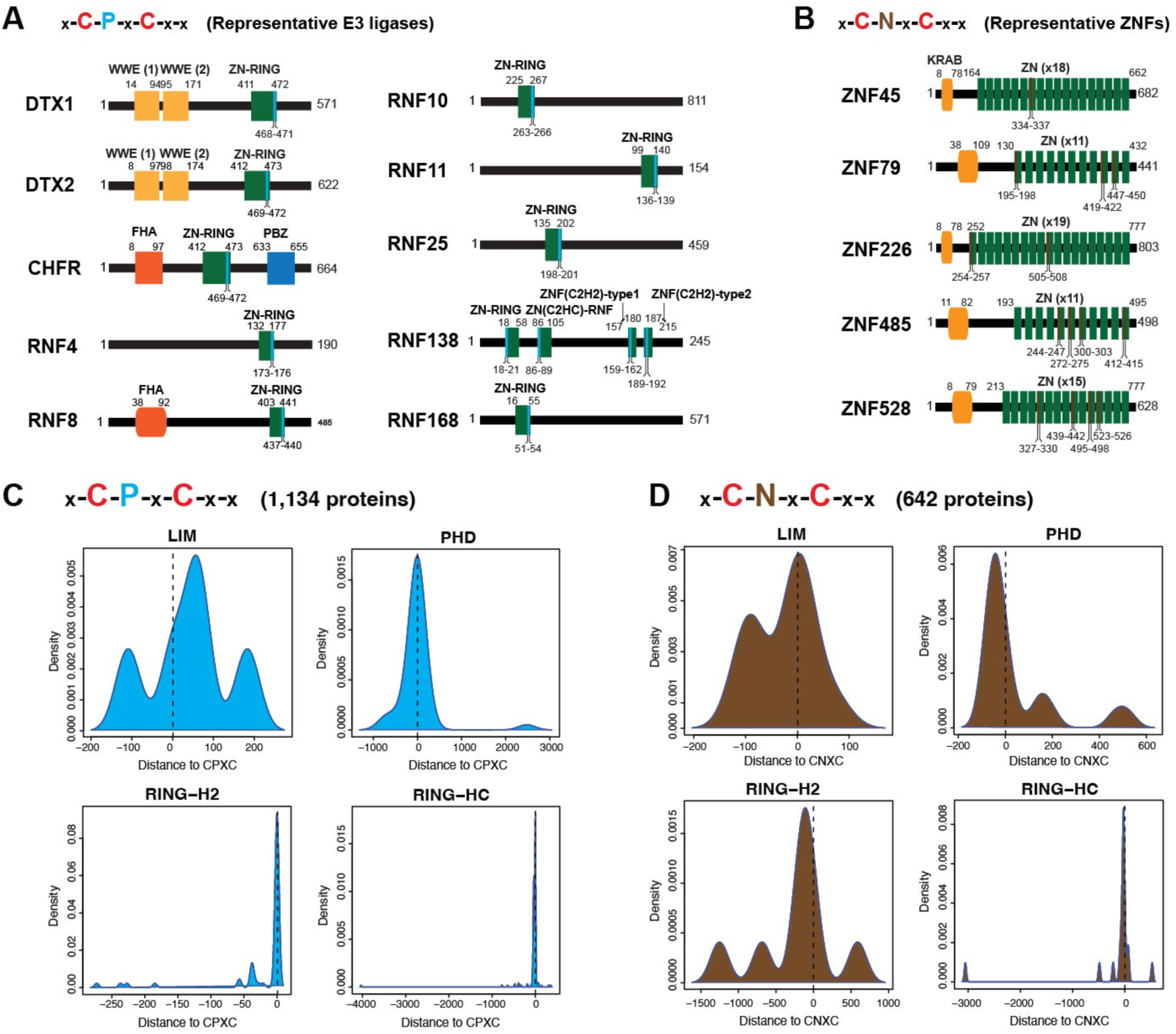
(A) Schematic representation of each CPxC’s localization in the representative E3 ligases. (B) Schematic representation of each CNxC’s localization in the representative ZNFs. Distance plot of each (C) CPxC and (D) CNxC motif to various domains: LIM, PHD, RING-H2, and RING-HC.

**Figure S5.**
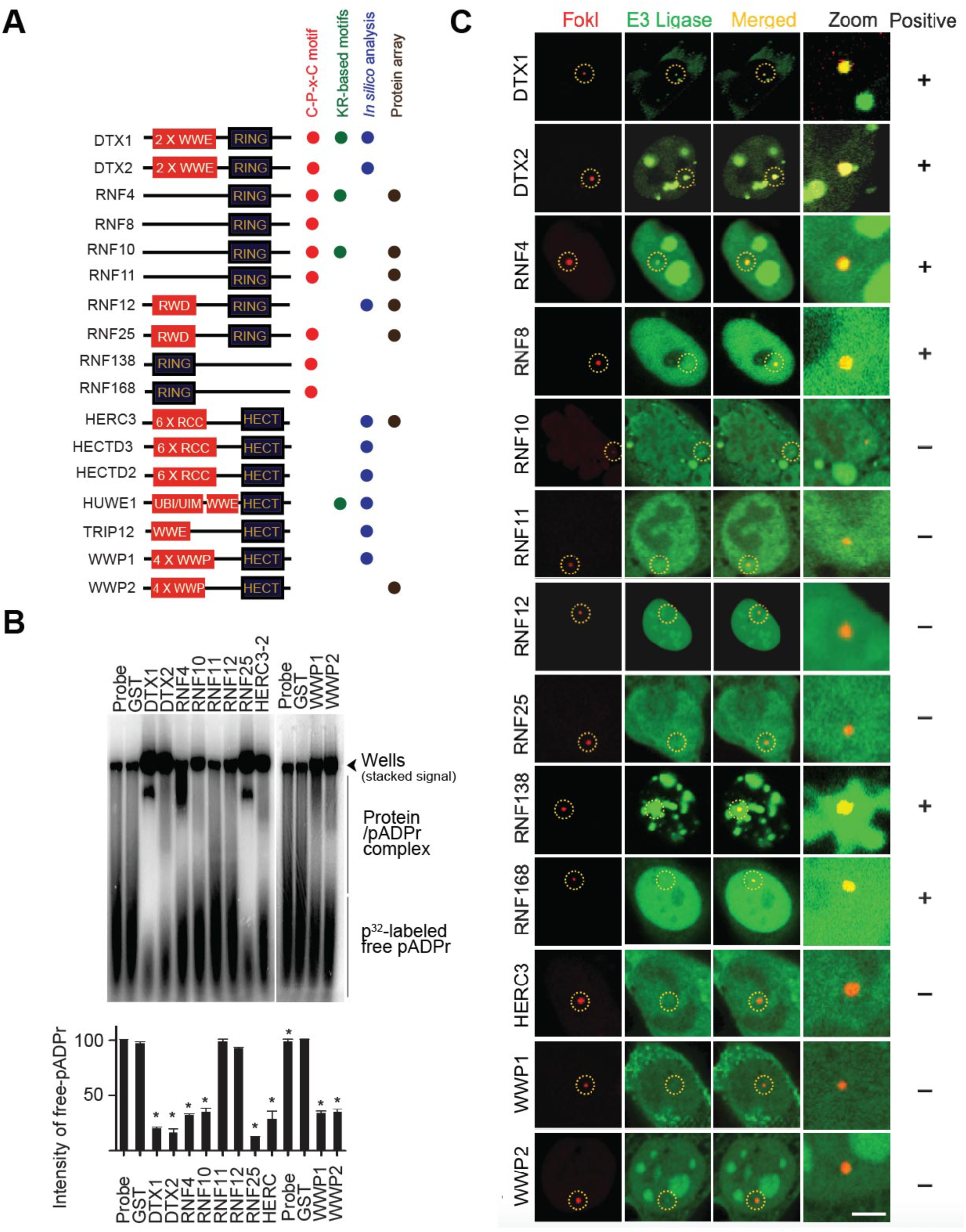
Screening of PAR-binding E3 ligases involved in DNA damage response (DDR) signaling. (A) Schematic illustrations of PAR-binding motifs in PAR-binding E3 ligases. The CPxC motif (Red) and KR-based motif (Green) in each PAR-binding E3 ligases are indicated. (B) A native electromobility gel shift assay for the validation of PAR-binding of newly predicted E3 ligases based on the new motifs. Intensity of each free [^32^P]-PAR was quantified and plotted. * P<0.05 by two tailed student t-test when compared GST binding to each of E3 ligases. *n.s.* not significant. (C) Screening of DDR E3 ligases. Co-localization of each GFP-fused PAR-binding E3s and mCherry–FokI nuclease at DSB site were represented. Scale bars, 10 μm.

**Figure S6.**
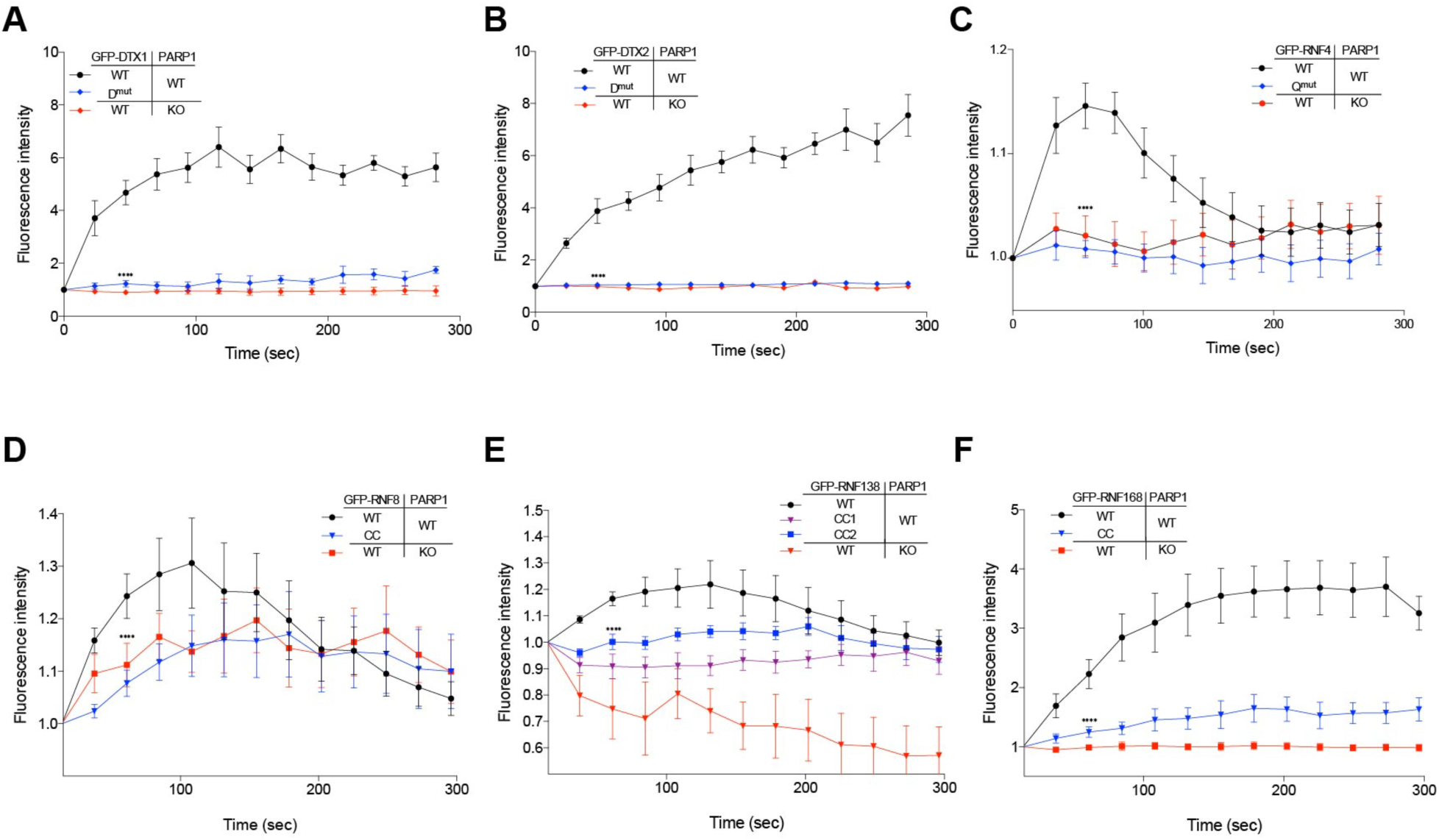
(A-F) GFP-fused wildtype and PBM E3s were expressed in either wildtype or PARP1 KO HeLa cells and microirradiated. Cumulative GFP signals at laser strips for (A) DTX1, (B) DTX2, (C) RNF4, (D) RNF8, (E) RNG138 and (F) RNF168 were measured and plotted., \*\*\*\**P*<0.0001, by one-way ANOVA when compared between indicated groups by Bonferroni’s posttest, Data represent mean ± *s.e.m.* from six independent cells (n=6).

**Figure S7.**
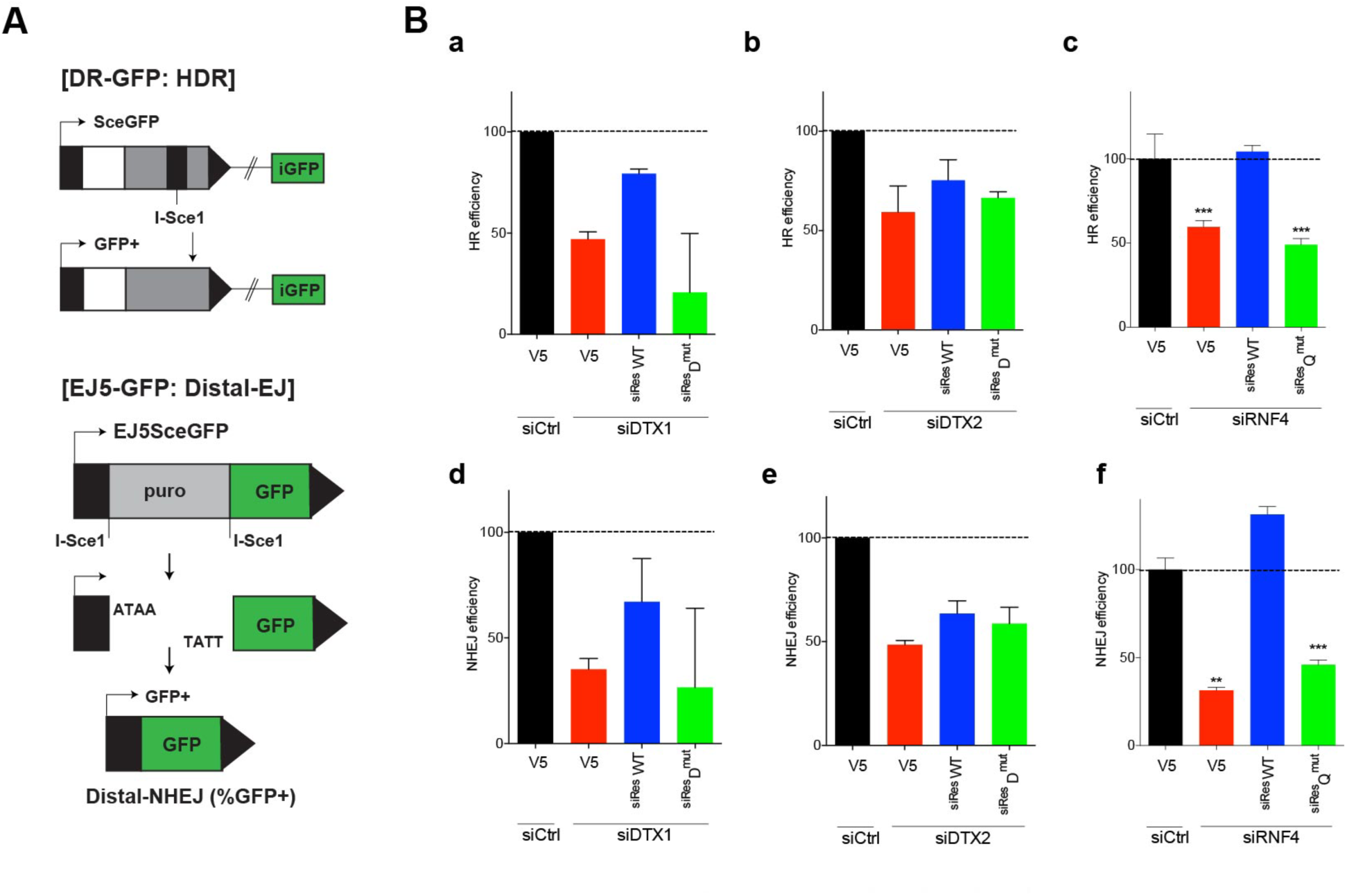
(A) Schematic illustrations of HR and NHEJ assay system used in this study. (Gunn and Stark, 2012) (Ba-Bf) DTX1, DTX2, and RNF4-targeted siRNA and siRNA-resistant constructs were transfected into HR- or NHEJ- reporter cell lines with I-SceI as indicated. After 72 h, GFP-positive cells were counted by FACS analysis. * P<0.001 by two tailed student t-test when compared control siRNA; V5 to siRNF4; V5 and siRNF4; V5 siResRNF4 Qmut.

### Supplementary Tables

**Table S1.** 356 PAR-binding proteins.

**Table S2.** Subcellular localization of 356 PAR-binding proteins.

**Table S3.** BioGRID analysis (Biological General Repository for Interaction Datasets).

**Table S4.** Classification of the most occurring sequence features using UniProt annotations.

**Table S5.** Overlapping protein list among data from 17k human proteome array, covalently PARylated proteins and non-covalent PAR-binding proteins.

**Table S6.** Overlapping domains and motifs among data from 17k human proteome array and four major screening methods.

**Table S7.** The classification of protein domains using 356 PAR-binding proteins.

**Table S8.** Overlapping protein list among data from experimentally identified using 17k human proteome array and two other theoretically predicted proteins.

**Table S9.** Characterization of two refined KR-based motifs.

**Table S10.** Overlapping protein list using approximately 20,000 human protein-coding genes among new motifs (CPxC/CNxC) and two refined KR motifs.

**Table S11.** Overlapping protein list using 515 human protein kinases among new motifs (CPxC/CNxC) and two refined KR motifs.

**Table S12.** Comparison of proteins containing new motifs (CPxC/CNxC).

**Table S13.** GO analysis using the gene list containing new motifs (CPxC/CNxC).

**Table S14.** Comparison of 614 human E3 ligases containing new motifs (CPxC/CNxC).

## Notes

### Competing Interest Statement

The authors have declared no competing interest.

